# Neutralizing effects of small molecule inhibitors and metal chelators on coagulopathic *Viperinae* snake venom toxins

**DOI:** 10.1101/2020.06.02.129643

**Authors:** Chunfang Xie, Laura Albulescu, Matyas A Bittenbinder, Govert Somsen, Freek Vonk, Nicholas R Casewell, Jeroen Kool

## Abstract

Animal-derived antivenoms are the only specific therapies currently available for the treatment of snake envenoming, but these products have a number of limitations associated with their efficacy, safety and affordability for use in tropical snakebite victims. Small molecule drugs and drug candidates are regarded as promising alternatives for filling the critical therapeutic gap between snake envenoming and effective treatment. In this study, by using an advanced analytical technique that combines chromatography, mass spectrometry and bioassaying, we investigated the effect of several small molecule inhibitors that target phospholipase A_2_ (varespladib) and snake venom metalloproteinase (marimastat, dimercaprol and DMPS) toxin families on inhibiting the activities of coagulopathic toxins found in *Viperinae* snake venoms. The venoms of *Echis carinatus, Echis ocellatus, Daboia russelii* and *Bitis arietans*, which are known for their potent coagulopathic toxicities, were fractionated in high resolution onto 384-well plates using liquid chromatography followed by coagulopathic bioassaying of the obtained fractions. Bioassay activities were correlated to parallel recorded mass spectrometric and proteomics data to assign the venom toxins responsible for coagulopathic activity and assess which of these toxins could be neutralized by the inhibitors under investigation. Our results showed that the phospholipase A_2_-inhibitor varespladib neutralized the vast majority of anticoagulation activities found across all of the tested snake venoms. Of the snake venom metalloproteinase inhibitors, marimastat demonstrated impressive neutralization of the procoagulation activities detected in all of the tested venoms, whereas dimercaprol and DMPS could only partially neutralize these activities at the doses tested. Our results provide additional support for the concept that combination of small molecules, particularly the combination of varespladib with marimastat, serve as a drug-repurposing opportunity to develop new broad-spectrum inhibitor-based therapies for snakebite envenoming.

## 1. Introduction

Bites by venomous cause 81,000 - 138,000 deaths per annum, with the majority occurring in the rural resource-poor regions of the tropics and sub-tropics [1]. The venomous snakes responsible for the vast majority of severe envenomings are members of the *Viperidae* and *Elapidae* families [2,3]. Elapid snakes have venoms that are highly abundant in neurotoxins that disable muscle contraction and cause neuromuscular paralysis [1,4]. Contrastingly, viper venoms typically contain numerous proteins that disrupt the functioning of the coagulation cascade, the hemostatic system, and tissue integrity [4,5]. Envenomings caused by these snakes can cause prominent local effects including necrosis, hemorrhage, edema and pain, and often result in permanent disabilities in survivors [6,7]. One of the most common but serious pathological effects of systemic viper envenoming is coagulopathy, which renders snakebite victims vulnerable to suffering lethal internal hemorrhages [8]. Venom induced coagulopathy presenting following bites by viperid snakes is thought to predominately be caused by venom enzymes such as phospholipase A_2_s (PLA_2_s), snake venom serine proteinases (SVSPs), and snake venom metalloproteinases (SVMPs) [9-11]. PLA_2_s can prevent blood clotting and induce anticoagulation by hydrolyzing phospholipids [12]. SVSPs can proteolytically degrade fibrinogen and can release bradykinins from plasma kininogens [13,14]. SVMPs work on various clotting factors and can degrade capillary basement membranes, thereby increasing vascular permeability and cause leakage [10,15,16]. These toxins can work synergistically causing local and systemic hemorrhage and coagulopathy.

The only specific therapies currently available for treating snake envenoming are animal-derived antivenoms. Consisting of immunoglobulins purified from hyperimmunized ovine or equine plasma/serum, these products save thousands of lives each year, but are associated with a number of therapeutics challenges, including limited cross-snake species efficacies, poor safety profiles and, for many snakebite victims residing in remote rural areas in developing countries, unacceptable issues with affordability and accessibility [17]. Small molecule toxin inhibitors are regarded as promising candidates for treating snakebite as they can seemingly generically block the enzymatic activities of venoms [18-20]. Varespladib, an indole-based nonspecific pan-secretory PLA_2_ inhibitor has been studied extensively for repurposing for snakebite. Having originally been investigated in Phase II and III clinical trials for treating septic shock, coronary heart disease and sickle cell disease-induced acute chest syndrome [21,22], varespladib has since been shown to be highly potent in suppressing venom-induced PLA_2_ activity both *in vitro* and *in vivo* in murine models [23]. Varespladib shows great promise against neurotoxic elapid snake venoms and has been shown to prevent lethality in murine *in vivo* models of envenoming [24] but is seemingly capable of also inhibiting certain myotoxic and coagulotoxic symptoms induced by snake venoms [25,26].

A number of other small molecules have shown promise for repurposing to inhibit SVMP venom toxins. Marimastat is a broad spectrum matrix metalloprotease inhibitor that functions by binding to the active site of matrix metalloproteinases where it coordinates the metal ion in the binding pocket [27,28]. As a water-soluble orally bioavailable matrix metalloproteinase inhibitor [29,30], marimastat reached phase II and III clinical trials for multiple solid tumor types [31-33], including pancreatic, lung, breast, colorectal, brain, and prostate cancer [34-36]. SVMPs are toxins which structurally and functionally are homologous to matrix metalloproteinses [37-39]. Like other compounds in this class of drugs (e.g. batimastat [40]), marimastat is a promising drug candidate for treating snakebite due to its inhibitory capabilities against SVMP toxins [41,42]. Dimercaprol, a historical drug approved by the World Health Organization (WHO) for treatment of heavy metal poisoning [43], contains two metal-chelating thiol groups and has long been used against arsenic, mercury, gold, lead and antimony intoxication [44-46]. It also represents a treatment option for Wilson’s disease in which the body retains copper. Moreover, it has been studied as a candidate for acrolein detoxification as it can effectively reduce the acrolein concentration *in vivo* in murine because of its ability to bind to both the carbon double bond and aldehyde group of acrolein. The water-soluble, tissue-permeable and licensed metal chelator, 2,3-dimercaptopropane-1-sulfonic acid (DMPS), is also suitable for treating acute and chronic heavy metal intoxication including lead, mercury, cadmium and copper [47,48]. It was recently shown that both dimercaprol and DMPS displayed potential for repurposing as small molecule chelators to treat snake envenoming [20], most probably by chelating and removing Zn^2+^ from the active site of Zn^2+^-dependent SVMPs. Of the two drugs, DMPS showed highly promising preclinical efficacy when used as an early oral intervention after envenoming by the SVMP-rich venom of the West African saw-scaled viper (*Echis ocellatus*), prior to later antivenom treatment with antivenom [20]. Thus, marimastat, dimercaprol and DMPS all represent promising candidates for drug repurposing as snakebite therapeutics, as they either inhibit SVMPs or chelate the Zn^2+^ ion required for SVMP catalysis.

In this paper, the coagulopathic properties of various snakes from the viper subfamily *Viperinae* (*Echis carinatus, E. ocellatus, Daboia russelii* and *Bitis arietans*) were evaluated using nanofractionation analytics in combination with a high-throughput coagulation assay, before the inhibitory capabilities of varespladib, marimastat, dimercaprol and DMPS against the coagulopathic toxicities of the resulting snake venom fractions were investigated. To this end, bioactivity chromatograms were acquired after fractionation, and parallel obtained mass spectrometry (MS) and proteomics data were used to correlate observed bioactivity with the identity of the venom toxins responsible for the observed enzymatic effects. Thus, we assessed the ability of varespladib, marimastat, dimercaprol and DMPS to neutralize the coagulopathic venom components. The results indicated that varespladib in combination with heavy metal chelators and/or broad-spectrum protease inhibitors could be viable first line therapeutic candidates for initial and adjunct treatment of coagulopathic snakebite envenoming.

## 2. Experimental

### 2.1 Chemicals

Water from a Milli-Q Plus system (Millipore, Amsterdam, The Netherlands) was used. Acetronitrile (ACN) and formic acid (FA) were supplied by Biosolve (Valkenswaard, The Netherlands). Calcium chloride (CaCl_2_, dehydrate, ≥ 99%) was from Sigma-Aldrich (Zwijndrecht, The Netherlands) and was used to de-citrate plasma to initiate coagulation in the coagulation assay. Phosphate buffered saline (PBS) was prepared by dissolving PBS tablets (Sigma-Aldrich) in water according to the manufacturer’s instructions and was stored at −4 °C for no longer than one week prior to use. Sodium citrated bovine plasma was obtained from Biowest (Nuaillé, France) as sterile filtered. The plasma (500 ml bottle) was defrosted in a warm water bath, and then quickly transferred to 15 ml CentriStar™ tubes (Corning Science, Reynosa, Mexico). These 15 ml tubes were then immediately re-frozen at −80 °C, where they were stored until use. Venoms were sourced from either wild-caught specimen maintained in or historical venom samples stored in the Herpetarium of the Liverpool School of Tropical Medicine. This facility and its protocols for the expert husbandry of snakes are approved and inspected by the UK Home Office and the LSTM and University of Liverpool Animal Welfare and Ethical Review Boards. The venom pools were from vipers with diverse geographical localities, namely: *B. arietans* (Nigeria), *D. russelii* (Sri Lanka), *E. carinatus* (India) and *E. ocellatus* (Nigeria). Note that the Indian *E. carinatus* venom was collected from a single specimen that was inadvertently imported to the UK via a boat shipment of stone, and then rehoused at LSTM on the request of the UK Royal Society for the Prevention of Cruelty to Animals (RSPCA). Venom solutions were prepared by dissolving lyophilized venoms into water to a concentration of 5.0 ± 0.1 mg/ml and were stored at −80 °C until use. The compounds varespladib (A-001), marimastat ((2S,3R)-N4-[(1S)-2,2-Dimethyl-1-[(methylamino)carbonyl] propyl]-N1,2-dihydroxy-3-(2-methylpropyl) butanedia-mide), dimercaprol (2,3-Dimercapto-1-propanol) and DMPS (2,3-dimercapto-1-propane-sulfonic acid sodium salt monohydrate) were purchased from Sigma-Aldrich. They were dissolved in DMSO (≥ 99.9%, Sigma-Aldrich) to a concentration of 10 mM and stored at −20 °C. Prior to use, these four compounds were diluted in PBS buffer to the described concentrations.

### 2.2 Venom nanofractionation

All venoms were nanofractionated onto transparent 384-well plates (F-bottom, rounded square well, polystyrene, without lid, clear, non-sterile; Greiner Bio One, Alphen aan den Rijn, The Netherlands) using a Shimadzu UPLC chromatography system (‘s Hertogenbosch, The Netherlands). The UPLC system was connected post-column to a modified Gilson 235P autosampler programmed for nanofractionation, which was controlled by the in-house written software Ariadne, or was post-column connected to a commercially available FractioMate™ nanofractionator (SPARK-Holland & VU, Netherlands, Emmen & Amsterdam) controlled by FractioMator software. The UPLC system was equipped with two Shimadzu LC-30AD parallel pumps, a Shimadzu SIL-30AC autosampler, a Shimadzu CTO-30A column oven, a Shimadzu SPD-M20A Prominence diode array detector and a DGU-20A5R Prominence degassing unit. All elements were remote controlled by the Shimadzu Lab Solutions software assisted by a Shimadzu CBM-20A System Controller. Venom solutions (5.0 ± 0.1 mg/ml) diluted in water to a concentration of 1.0 mg/ml were injected (50 μl) for nanofractionation after gradient liquid chromatography (LC). A Waters XBridge reverse-phase C18 column (250 × 4.6 mm with 3.5-μm pore-size particles) and a Shimadzu CTO-30A column oven maintained at 30 °C were used for LC separations. The total eluent flow rate was 0.5 ml/min and was controlled by the two Shimadzu LC-30AD parallel pumps. The gradient separation was carried out by linearly increasing mobile phase B from 0 to 50% during the first 20 min, from 50% to 90% during the following 4 min, and was then kept at 90% for 5 min. Subsequently, mobile phase B was decreased from 90 to 0% in 1 min and kept at 0% for 10 min. Mobile phases A consisted of 98% H_2_O, 2% ACN and 0.1% FA, while mobile phase B consisted of 98% ACN, 2% H_2_O and 0.1% FA. A 9:1 (v/v) split of the column effluent was applied, of which the smaller fraction was sent to the UV detector followed by MS, and the larger fraction was directed to the nanofraction collector. The nanofractionator was set to continuously collect fractions of 6 s/well. After fraction collection, the transparent 384-well plates were freeze-dried overnight using a Christ Rotational Vacuum Concentrator (RVC 2−33 CD plus, Zalm en Kipp, Breukelen, The Netherlands) equipped with a cooling trap operated at −80 °C. The freeze-dried plates were stored at −20 °C until the bioassays were performed.

### 2.3 Plasma coagulation activity assay

The HTS plasma coagulation assay used in this study was developed by Still *et al.* [49]. CaCl_2_ was dissolved in water to a concentration of 20 mM at room temperature. A 15 ml CentriStar™ tube with frozen plasma was defrosted to room temperature in a warm water bath and then centrifuged at 2000 rpm (805 × g) for 4 min to remove potential particulate matter. Stock solutions (10 mM) of the compounds under investigation (i.e. varespladib, marimastat, dimercaprol and DMPS) were diluted in PBS buffer to the required concentrations. Of these diluted solutions, 10 μl were pipetted into all plate wells containing freeze-dried venom fractions by using a VWR Multichannel Electronic Pipet (10 μl of PBS were used for venom-only analyses as a control). Next, plates were centrifuged for 1 min at 2000 rpm (805 × g) in a 5810 R centrifuge (Eppendorf, Germany) and then pre-incubated for 30 min at room temperature. The final concentrations of the inhibitor solutions used in the coagulation bioassay were 20 μM, 4 μM and 0.8 μM, and in some cases 0.16 μM, 0.032 μM and 0.0064 μM.

Following incubation, 20 μl of the CaCl_2_ solution was pipetted into each well of a 384-well plate with vacuum-centrifuged (to dryness) venom fractions, followed by 20 μl of centrifuged plasma using a Multidrop™ 384 Reagent Dispenser (Thermo Fisher Scientific, Ermelo, The Netherlands) after in-between rinsing the Multidrop with Milli-Q. Immediately after plasma addition, the plate was placed in a Varioskan™ Flash Multimode Reader (Thermo Fisher Scientific, Ermelo, The Netherlands) and a kinetic absorbance measurement was performed at a wavelength of 595 nm at room temperature for 100 min. All analyses were performed at least in duplicate. The slope of the signal obtained for each well was normalized by dividing the slope to the median of all the slope signals from all wells in that measurement. The coagulation curves were plotted versus the chromatographic retention time for each fraction collected in three different ways (very fast coagulation activity, slightly/medium increased coagulation activity and anticoagulation activity) to fully depict both the procoagulation and anticoagulation activities in each well. Detailed explanations on the rationale of processing and plotting the data in this way is provided by Slagboom *et al*. and Xie *et al*. [50,51], and can be found in the Supporting Information (Section S1).

### 2.4 Correlation of biological data with MS data

The corresponding accurate mass(es) and proteomics data for each venom fraction in this study have already been acquired by Slagboom *et al.* [51] and as such were correlated with the bioactivity chromatograms obtained in the current study. For venoms under study in this research that were not studied by Slagboom *et al.* [51], the same procedure as described by Slagboom *et al.* [51] was followed to acquire and process proteomics data on these snake venoms. The UniprotKB database was used to determine the toxin class and any known functions for the relevant toxins thought to be responsible for the observed coagulopathic toxicities. For LC separations performed at different times and in different labs, the retention times of eluting snake venom toxins may differ slightly. The LC-UV chromatograms (measured at 220 nm, 254 nm, and 280 nm), which provided characteristic fingerprint profiles for each venom fraction, were used to negotiate these retention time shifts. By using the LC-UV data, the chromatographic bioassay data from this study was correlated with the MS total-ion currents (TICs), extracted-ion chromatograms (XICs), and proteomics data obtained by Slagboom *et al.* [51]. In order to construct useful XICs, MS spectra were extracted from the time frames that correlated with regions in the chromatograms for each bioactive peak. Then, for all *m/z* values showing a significant signal observed in the mass spectra, XICs were plotted. In turn, these XICs were used for matching with peak retention times of bioactive compounds in the chromatograms. The exact masses matching the bioactives were tentatively assigned based on matching peak shape and correlation with retention times in bioassay traces. More specifically, the *m/z*-values in the MS data were correlated to each bioactive peak using the accurate monoisotopic masses determined by applying the deconvolution option in the MS software. For the proteomics data, in-well tryptic digestions were performed by Slagboom *et al.* [51] on snake venom fractions. These proteomics results could directly be correlated to coagulopathic activity that was indicated by the bioassay chromatograms.

## 3. Results

In this study, a nanofractionation approach was used to evaluate the inhibitory effects of varespladib, marimastat, dimercaprol and DMPS on the coagulopathic properties of venom toxins fractionated from a variety of *Viperinae* snake species. A recently developed low-volume HTS coagulation bioassay was used to assess the coagulation activities of LC-fractionated venoms in a 384-well plate format. These coagulopathic activities were correlated to parallel obtained MS and proteomics data to determine which specific venom toxins were neutralized by the potential inhibitors. All analyses were performed at least in duplicate using venom concentrations of 1.0 mg/ml.

### 3.1 Inhibitory effects of varespladib, marimastat, dimercaprol and DMPS on Echis venoms

Various geographically distinct saw-scaled viper venoms (genus *Echis*) were investigated in this study, specifically from the Indian species *E. carinatus* and the west African species *E. ocellatus*. The inhibitory effects of varespladib, marimastat, dimercaprol and DMPS against the coagulopathic activities observed for LC fractions of both venoms were investigated in a concentration-dependent fashion (Figures 1-2). Duplicate bioassay chromatograms together with a detailed description of each coagulopathic peak observed are presented in the Supporting Information (Section S1).

**Figure 1.**
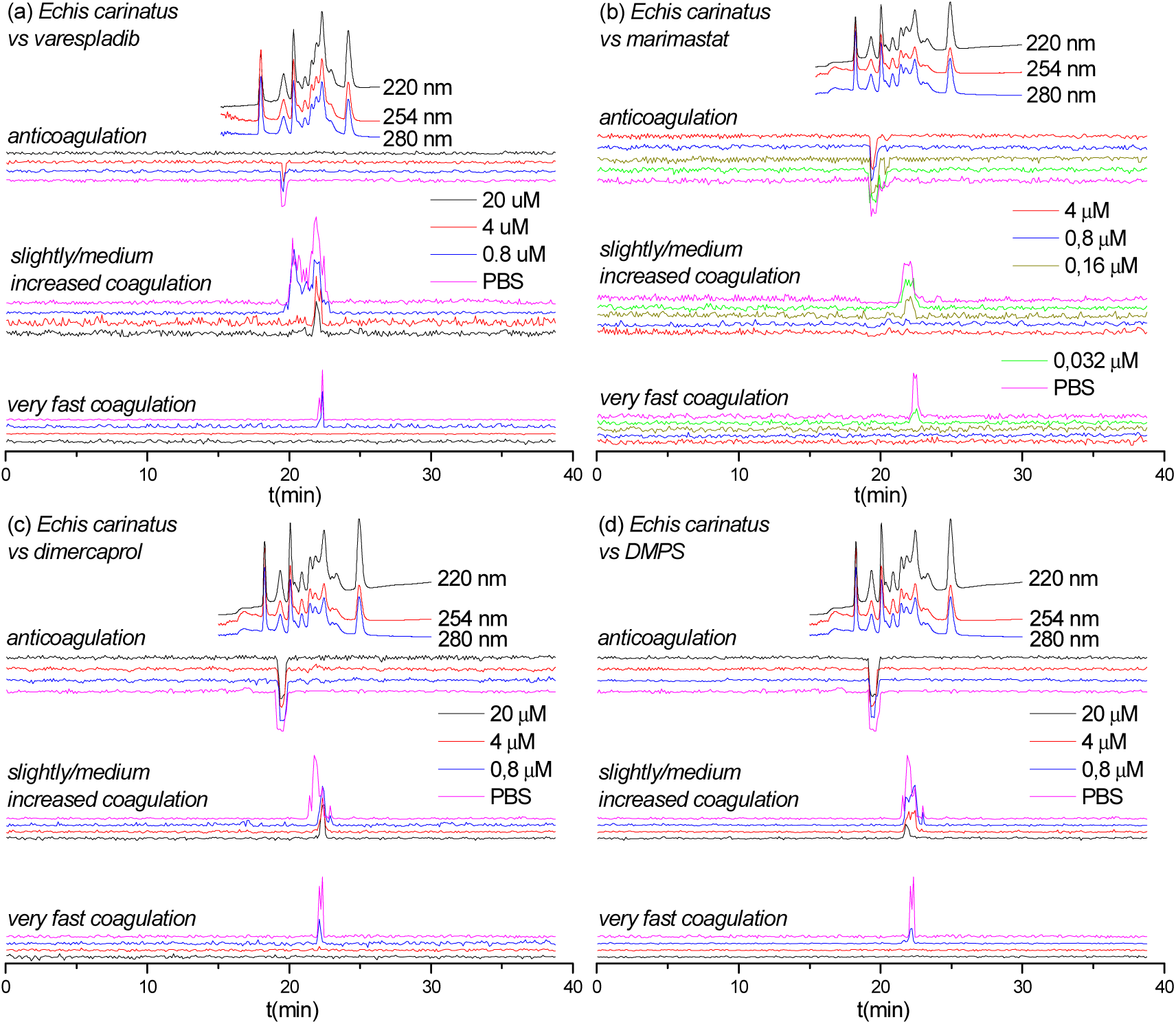
UV absorbance chromatograms and reconstructed coagulopathic toxicity chromatograms of nanofractionated venom toxins from *E. carinatus* venom in the presence of different concentrations of (a) varespladib, (b) marimastat, (c) dimercaprol and (d) DMPS.

**Figure 2.**
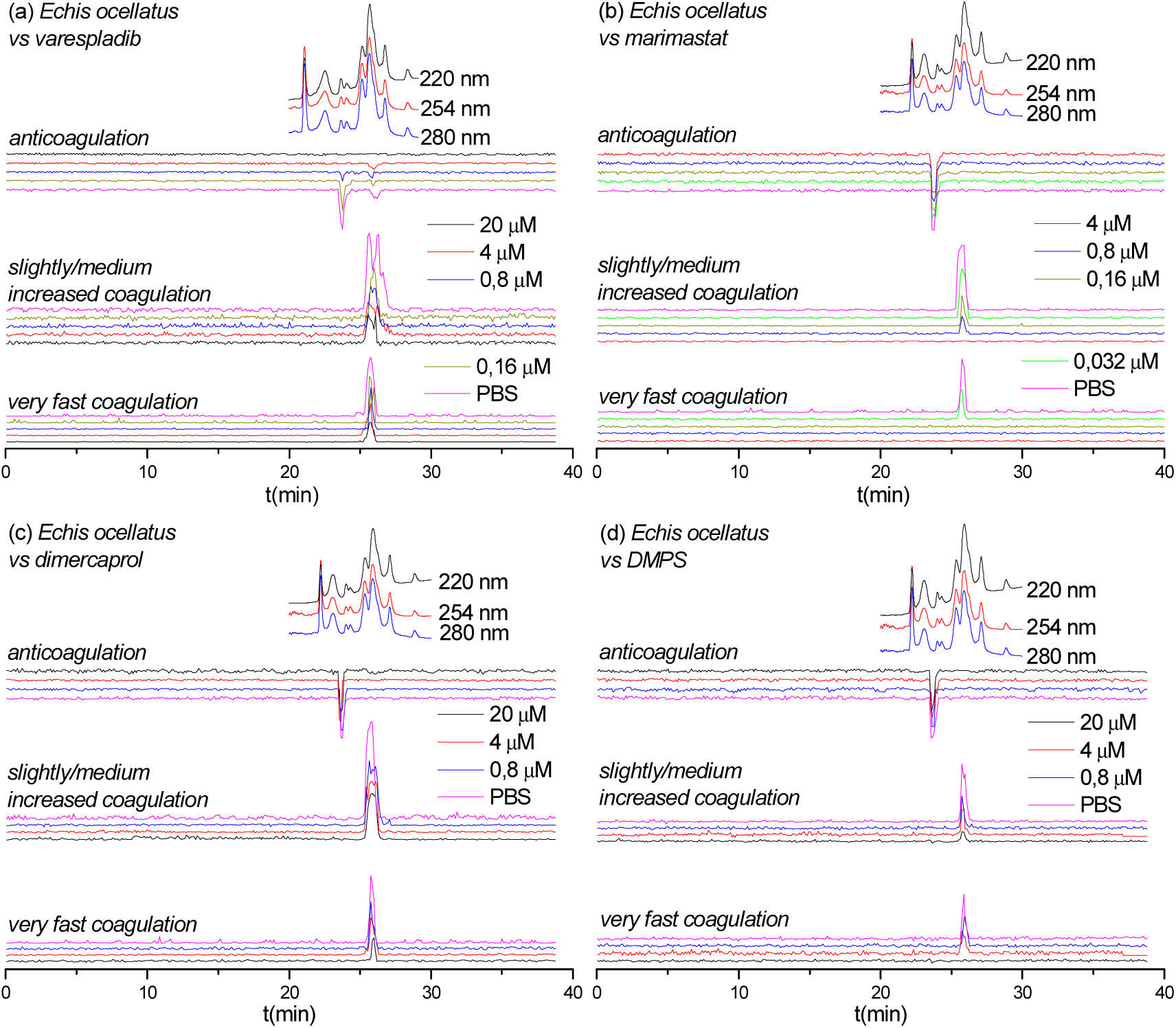
UV absorbance chromatograms reconstructed coagulopathic toxicity chromatograms of nanofractionated venom toxins from *E. ocellatus* venom in the presence of different concentrations of (a) varespladib, (b) marimastat, (c) dimercaprol and (d) DMPS.

Figure 1 shows the bioassay chromatograms of nanofractionated venom toxins from *E. carinatus* in the presence of different concentrations of varespladib, marimastat, dimercaprol and DMPS. In the venom-only analysis, potent procoagulation activities were observed in the very fast coagulation chromatogram (22.0-22.9 min) and the slightly/medium increased coagulation chromatogram (21.2-23.1 min and/or 19.9-21.2 min), while anticoagulation activities were observed in the anticoagulation chromatogram (19.1-19.9 min). Interestingly, the PLA_2_-inhibitor varespladib inhibited both the anticoagulation and procoagulation activities, with the exception of one major peak observed in the slightly/medium increased coagulation chromatogram. In contrast, marimastat, dimercaprol and DMPS only exerted inhibitory effects on the procoagulation activities of *E. carinatus* venom. The anticoagulation activity of *E. carinatus* venom was fully inhibited by varespladib at a 20 μM concentration, while the very fast procoagulation activity was fully inhibited by varespladib, dimercaprol and DMPS at a concentration of 4 μM. Marimastat superseded the other small molecules by fully inhibiting the very fast procoagulation activity at a concentration of 0.16 μM. The slightly/medium increased coagulation activity was fully inhibited by 0.8 μM marimastat, but a sharp positive peak (21.7-22.2 min) was still retained following incubation with 20 μM varespladib. Dimercaprol only inhibited the front peak (21.3-22.1 min) present in the slightly/medium increased coagulation activity chromatogram, while DMPS inhibited mostly the tailing part (22.2-23.1 min) of this peak at its highest concentration tested (20 μM). Overall, DMPS was found to be more effective than dimercaprol in abrogating the procoagulation toxicities of *E. carinatus* venom. These findings demonstrate that the tested inhibitors have different specificities, but that marimastat most effectively inhibits procoagulant components and varespladib anticoagulant components of *E. carinatus* venom.

Figure 2 shows the bioassay chromatograms for nanofractionated toxins from *E. ocellatus* venom in the presence of different concentrations of varespladib, marimastat, dimercaprol and DMPS. In the venom-only analysis, we observed similar results to those obtained for *E. carinatus* venom; multiple co-eluting sharp peaks were present in the very fast coagulation chromatogram (25.1-26.2 min), the slightly/medium increased coagulation chromatogram (25.1-27.1 min) and the anticoagulation chromatogram (23.4-24.4 min). All peaks decreased in height and width with increasing varespladib concentrations. The potent negative peak (23.4-24.4 min) in the anticoagulation chromatograms was fully inhibited by 4 μM varespladib and the later eluting weakly negative peak (25.9 min) by 20 μM varespladib. While full inhibition of anticoagulation activities was achieved, the procoagulation activities were not fully inactivated at the highest varespladib concentration tested (20 μM). However, both the very fast coagulation activity and the slightly/medium increased coagulation activity were also somewhat reduced by varespladib in a concentration-dependent fashion. Similar findings, whereby both very fast and slightly/medium increased coagulation were reduced in a concentration dependent manner but not fully abrogated, were also observed for dimercaprol, although this inhibitor had no effect on anticoagulant venom activities. Marimastat and DMPS also had no effect on anticoagulant venom activity, but effectively inhibited the procoagulant actions of *E. ocellatus* venom. Very fast procoagulation activity was fully inhibited at a lower concentration of marimastat (0.16 μM) than DMPS (20 μM), while slightly/medium increased coagulation activity was fully inhibited by 4 μM marimastat compared with almost complete inhibition observed when using 20 μM DMPS. Thus, similar to findings with *E. carinatus*, marimastat exhibited superior inhibition of procoagulant venom activities, while varespladib was the only inhibitor capable of abrogating anticoagulant venom effects.

### 3.2 Inhibitory effect of varespladib, marimastat, dimercaprol and DMPS on Daboia russelii venom

Next, we assessed the inhibitory capability of the same small molecule toxin inhibitors on a *Viperinae* snake from a different genus – the Russell’s viper (*Daboia russelii*), which is highly medically-important species found in south Asia [52-54]. The inhibitory effects of varespladib, marimastat, dimercaprol and DMPS on the venom of *D. russelii* are shown in Figure 3. Duplicate bioassay chromatograms for the *D. russelii* venom analyses can be found in the Supporting Information in Section 2. For the venom-only analysis, a strong positive peak was observed for both the very fast coagulation activity (21.5-22.4 min) and for the slightly/medium increased coagulation activity (21.5-22.8 min). A very broad and strong negative activity peak (18.6-21.5 min) was also observed, demonstrating potent anticoagulation activity. In terms of procoagulant venom effects, both very fast and slightly/medium increased coagulation activities decreased dose-dependently in the presence of varespladib, marimastat and dimercaprol, although neither varespladib nor dimercaprol could fully neutralize these activities. However, in line with the earlier findings for *Echis* spp, full neutralization of both types of procoagulation were observed with marimastat, at 0.8 μM for very fast coagulation activity and at 4 μM for slightly/medium increased coagulation activity. As anticipated, and again in line with findings observed with *Echis* spp., neither of the SVMP-inhibitors (marimastat and dimercaprol) abrogated anticoagulant venom activity. In contrast, varespladib showed potent inhibition of anticoagulation, as the broad and potent negative peak (18.6-21.5 min) decreased to only a very minor negative peak (19.5-20.2 min; 20 μM varespladib) with increasing varespladib concentrations. DMPS showed no inhibition on both the procoagulant and anticoagulant venom activities at tested concentrations of 20 μM and 4 μM on *D. russelii* venom (Figure 3 (d)).

**Figure 3.**
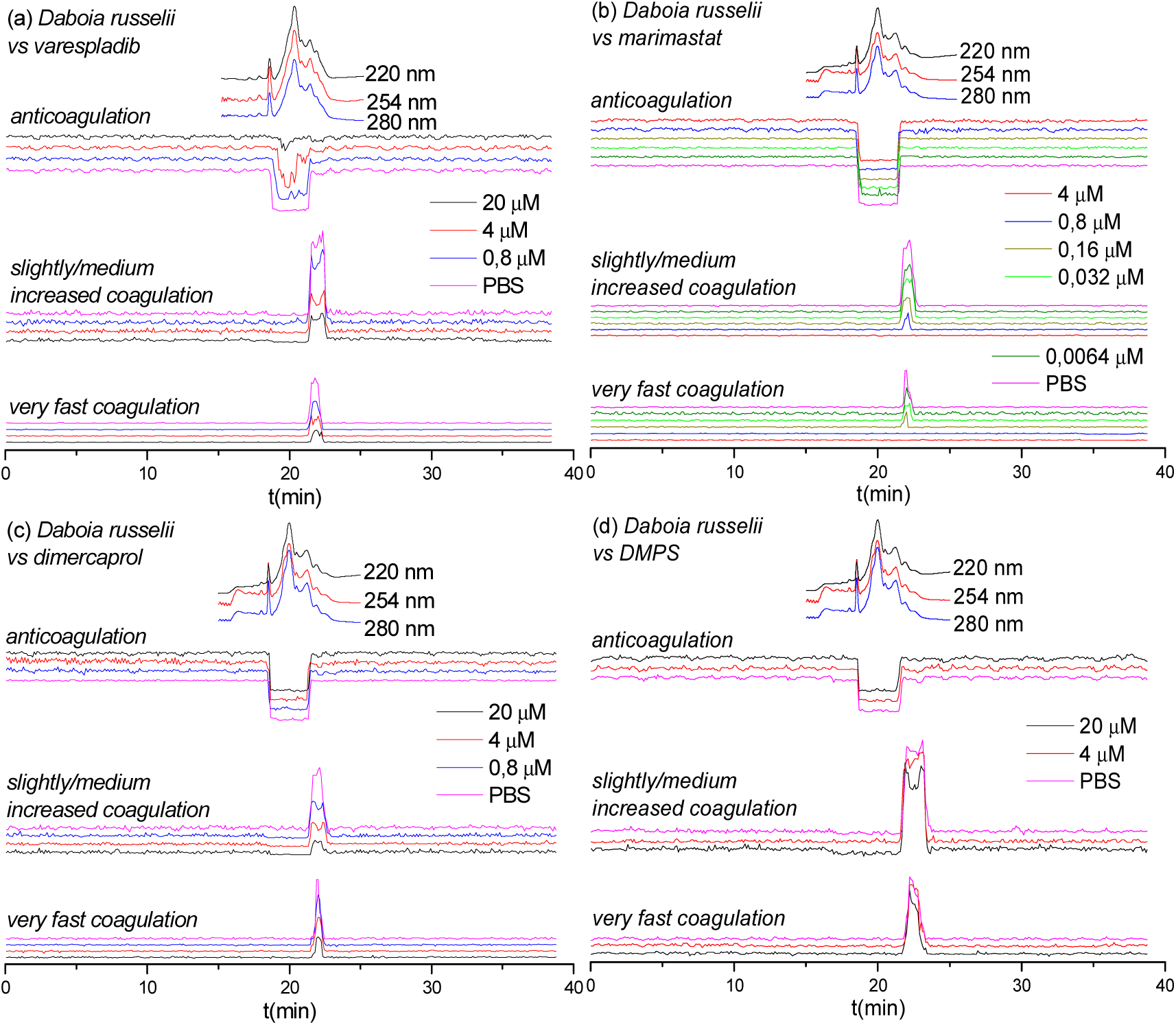
UV absorbance chromatograms and reconstructed coagulopathic toxicity chromatograms of nanofractionated venom toxins from *D. russelii* venom in the presence of different concentrations of (a) varespladib, (b) marimastat, (c) dimercaprol and (d) DMPS.

### 3.3 Inhibitory effects of varespladib and marimastat on Bitis arietans venom

The inhibitory effects of varespladib and marimastat on the coagulopathic properties of venom of the puff adder (*B. arietans*) which is found widely distributed across sub-Saharan Africa and parts of the Middle East, are shown in Figure 4. Duplicate bioassay chromatograms for the *B. arietans* venom analyses are shown in the Supporting Information in Section 3. In the venom-only analyses, anticoagulation activity was observed as two sharp negative peaks in the bioactivity chromatograms (16.2-16.7 min and 16.7-17.1 min), however no procoagulation activity was detected, which is consistent with previous findings using this venom [55]. Consequently, of the three SVMP-inhibitors used elsewhere in this study, we only selected marimastat for assessment of toxin inhibition as a control for the PLA_2_-inhibitor varespladib. In line with findings from the other *Viperinae* species under study, increasing concentrations of varespladib resulted in full inhibition of the two negative anticoagulation peaks, at concentrations of 0.16 μM and 0.8 μM, respectively. Conversely, and also in line with our earlier findings, no inhibitory effects were observed with marimastat, even at concentrations of 20 μM.

**Figure 4.**
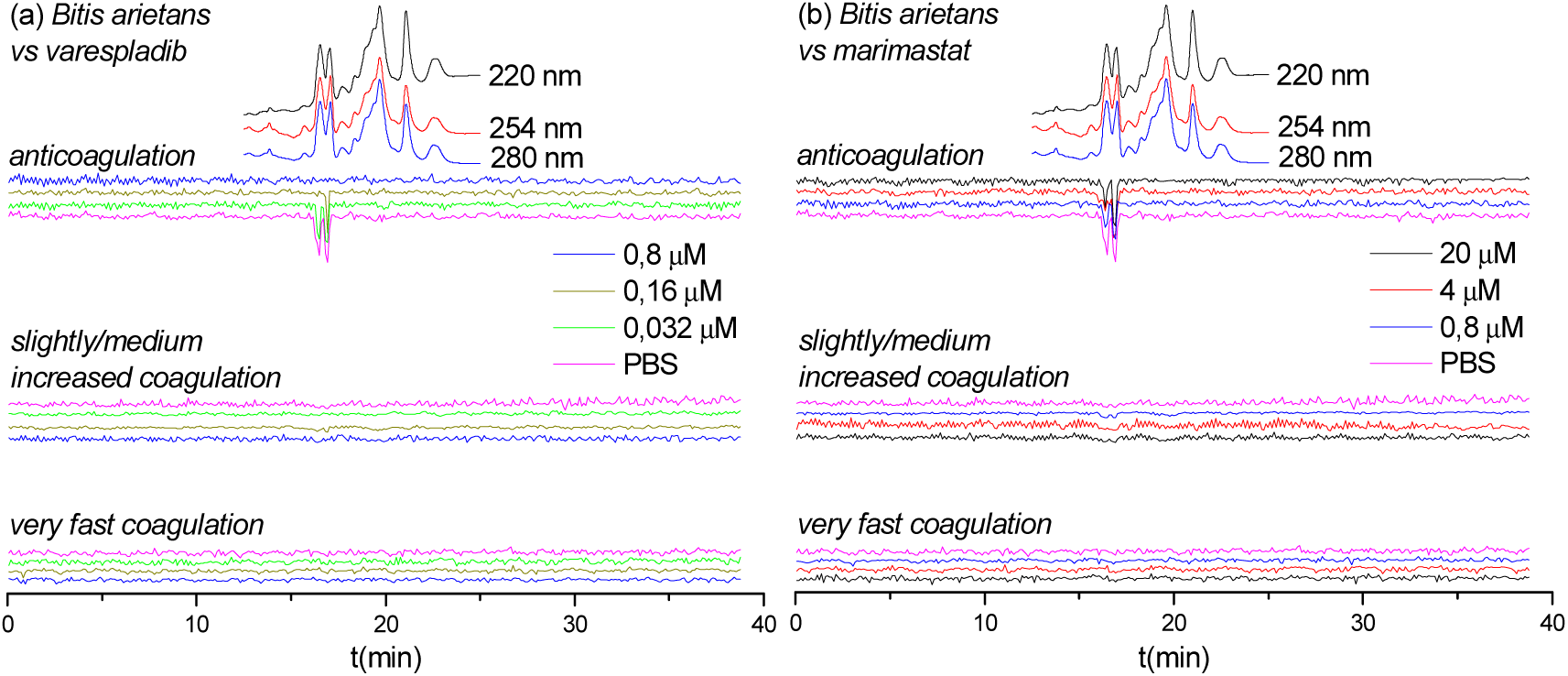
UV absorbance chromatograms and reconstructed coagulopathic toxicity chromatograms of nanofractionated venom toxins from *B. arietans* venom in the presence of different concentrations of (a) varespladib and (b) marimastat.

### 3.4 Identification of coagulopathic venom toxins neutralized by varespladib, marimastat, dimercaprol and DMPS

MS and proteomics data previously obtained by Slagboom *et al.* [51] was used to assign the venom toxins responsible for the observed coagulation activities are listed in Table 1. All tentatively identified anticoagulant PLA_2_s are provided in Table 1, including those found in our study not previously described as possessing anticoagulant properties in the UniprotKB database. For those toxins for which no exact mass data could be acquired by LC-MS, only the proteomics mass data retrieved from Mascot searches are presented.

**Table 1.**
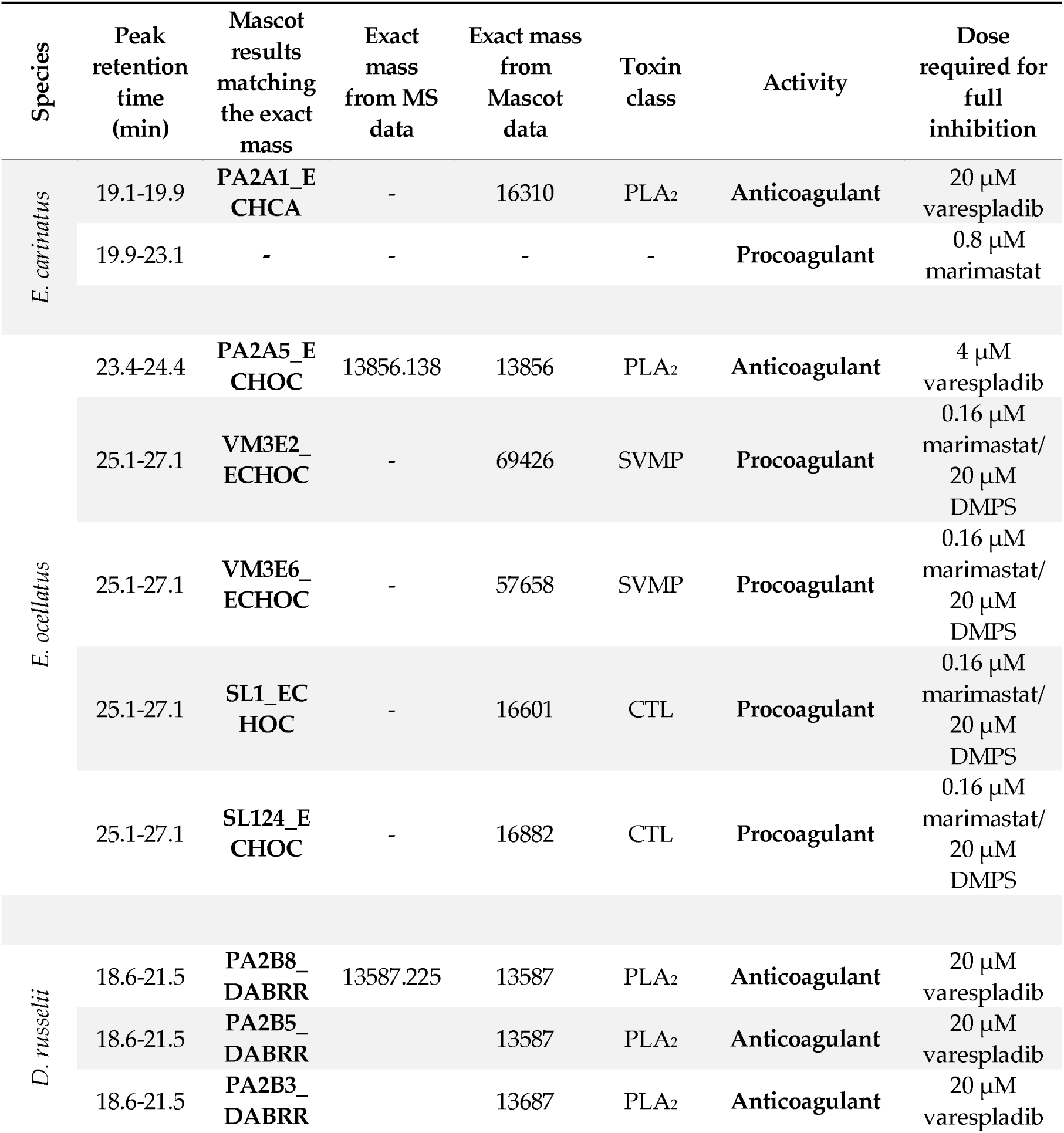
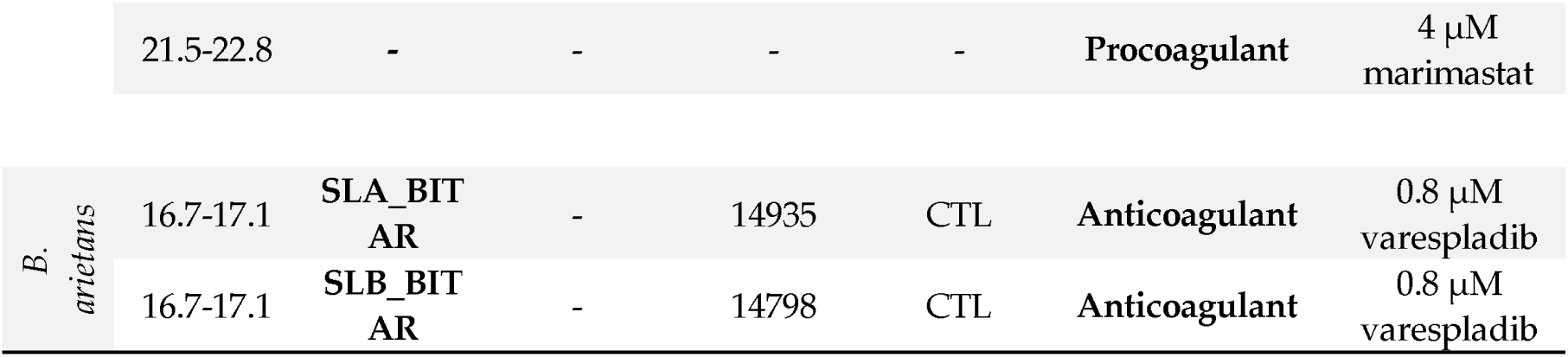
Correlated MS and proteomics data for associated coagulopathic venom toxins. (Peak retention times are adapted from Figures 1-4; PLA_2_ = phospholipase A_2_; SVMP = Snake Venom Metalloproteinase; CTL = C-Type Lectin).

Based on results from Figures 1-4 and Table 1, the inhibitory effects of varespladib, marimastat, dimercaprol and DMPS on individual *Viperinae* venom toxins were assessed. PLA_2_ toxins were identified as components responsible for anticoagulation in all species studied, except for *B. arietans*, for which CTLs were identified. All these identified anticoagulant toxins were fully abrogated by varespladib at various concentrations, as indicated in Table 1. The CTLs identified for *B. arietans* venom highly likely co-eluted with anticoagulants. However, no other anticoagulants were identified from this venom. No procoagulant toxins could be identified for the procoagulant peaks from the Mascot results for *E. carinatus, D. russelii* and *B. arietans* venoms. The procoagulants identified for *E. ocellatus* venom were both SVMPs and CTLs. All these toxins could be fully abrogated by marimastat at 0.16 μM or by DMPS at 20 μM. The procoagulant activity detected for *E. carinatus* and *D. russelii* venoms could be fully inhibited by marimastat at low concentrations. It was reasonable to speculate that the procoagulant toxins responsible for these activities were mainly SVMPs. In the cases where multiple toxins elute closely, unambiguously assigning single toxins to each detected bioactivity is difficult. For bioactive compounds that eluted in activity peaks which were only partly inhibited, it was difficult to critically determine which of them was abrogated. This would require further improving LC separations under toxin non-denaturating and MS compatible eluent conditions. As critical note it has to be mentioned that despite venom toxins generally are rather stable, during chromatography within the nanofractionation analytics pipeline some venom toxins might have (partly) denatured and thereby lost their activity. A detailed description of the results discussed here is provided in the Supporting Information in Section S4.

## 4. Discussion

The results of this study show that the chosen four small molecule inhibitors (i.e. varespladib, marimastat, dimercaprol and DMPS) are capable of inhibiting, albeit to different extents and specificities, the coagulopathic activities of individual toxins present in *Viperinae* venoms. While this is consistent with previous work on small molecule inhibitors and chelators exhibiting anti-hemorrhagic and anti-procoagulant activities of snake venoms [9,20,23,41,42,56], here we have detailed the relative specificities of these molecules. Our findings reveal that not only is varespladib effective against the activity of anti-coagulant PLA_2_ toxins, but also shows some inhibitory activity against procoagulant venom toxins. Contrastingly, of the SVMP-inhibitors tested, we demonstrate that their specificities are restricted to effects on procoagulant venoms toxins, and that the peptidomimetic hydroxamate inhibitor marimastat outperforms the metal chelators DMPS and dimercaprol in terms of potency.

There is an urgent need for stable, economical and effective snakebite treatments that can be administered in the field and in rural areas where medical access is limited. Small molecule inhibitors that specifically target a number of key classes of snake venom toxins have recently gained interest as candidates for therapeutic alternatives to antivenom, either as solo therapies or in toxin-specific inhibitor combinations, or as adjunctive treatment in combination with conventional antivenoms [18]. These compounds show great promise for the long-term development of affordable, broad-spectrum, first-aid and clinical treatment of tropical snakebite for a variety of reasons. The advantages of repurposing licensed medicines (e.g. DMPS, dimercaprol) or at least phase II-approved drug candidates (e.g. marimastat, varespladib) for snakebite is that these molecules have demonstrated safety profiles, and thus development could be significantly shortened as these agents have extensive pharmacokinetic, bioavailability and tolerance data already associated with them [23,57,58]. The small size of these compounds, compared with conventional antibodies, confer desirable drug-favourable properties enabling rapid and effective tissue penetration and, depending on the pharmacokinetics and physicochemical properties of specific inhibitors, often make them amenable for oral delivery [57,59,60]. Indeed, both varespladib and DMPS have already been demonstrated to confer preclinical efficacy against snakebite via the oral route [20,59,60]. Also, because these inhibitors can be produced in large amounts using efficient, low cost and validated synthetic procedures, they are promising candidates as more affordable treatments than conventional antivenoms for snakebite victims found in low- and middle-income countries [20,61].

In addition to these desirable characteristics, certain small molecule inhibitors have demonstrated broad inhibition of specific toxin families across diverse medically-important snake species [23,55] - as also evidenced here for the various coagulopathic toxins found across *Viperinae* venoms. Thus, one of the great strengths of these inhibitors lies in their ability to neutralize the activities of multiple toxin isoforms across different snake species by exploiting the similarities of the catalytic sites of action found within specific toxin families [58]. Thus, small molecule inhibitors show potential to circumvent the therapeutic challenge that venom variation presents. However, these compounds typically target only a single family of enzymatic toxins although varespladib seems capable of targeting more than one family (see Figures 1(a), 2(a), 3(a)), thus presenting a challenge for these molecules to become standalone therapeutics, as other non-inhibited toxins seem likely to still cause pathology in snakebite victims. It is therefore more likely that small molecule inhibitors will need to be combined into therapeutic mixtures, either with other toxin inhibitors or monoclonal antibodies, to generate snakebite therapeutics capable of neutralizing a wide breadth of snake venoms [26,57,58]. We recently described such an approach, whereby combining the PLA_2_-inhibitor varespladib and the SVMP-inhibitor marimastat resulted in a therapeutic combination capable of preclinically neutralizing lethality caused by the venom of diverse haemotoxic viper species from Africa, south Asia and central America [55]. Our findings in this study provide complementary data supporting the future translation of this therapeutic mixture, since we find here that varespladib effectively inhibits anticoagulant venom toxins, while marimastat potently inhibits procoagulant venom toxins (Figures 1(b), 2(b), 3(b)). On the other hand, small molecule inhibitors could serve as valuable prehospital snakebite treatments to delay the onset of severe envenoming before the arrival of victims to secondary or tertiary healthcare facilities to receive subsequent therapy (i.e. conventional antivenoms). Indeed, compounds such as varespladib and DMPS are already being explored in this regard [20,59,60], as they represent promising candidates to be used as bridging therapies for delaying the major effects of envenomation, and reducing the long time it typically takes rural, isolated, impoverished snakebite victims to receive any form of treatment. This is important, because treatment delays are known to have major detrimental impacts on patient outcomes following snakebite [62,63].

## 5. Conclusions

In this study, a recently developed HTS coagulation assay was combined with LC fractionation and parallel obtained MS and proteomics data to assess the neutralizing potency of several small molecule inhibitors and chelators (i.e. varespladib, marimastat, dimercaprol and DMPS) against the coagulopathic activities of individual toxins found in the venoms of *Viperinae* snakes. Both procoagulant and anticoagulant activities were detected in *E. carinatus, E. ocellatus* and *D. russelii* venoms, while only anticoagulant activity was present in *B. arietans* venoms (at 1.0 mg/ml venom injected). The PLA_2_-inhibitor varespladib potently inhibited the anticoagulant activities detected in all venoms, except for *D. russelii*, for which almost complete inhibition was observed. In addition, and surprisingly, varespladib showed some degree of inhibition against procoagulant venom activities across the various venoms, despite these activities not being mediated by PLA_2_ toxins. Marimastat potently inhibited procoagulant activities across the venoms tested, but was unsurprisingly ineffective against anticoagulant venom activities. These findings are consistent with the anticipated mechanism of action underlying the inhibitory activity of marimastat - specifically binding to the active site of, often procoagulant, snake venom metalloproteinases. Marimastat outperformed dimercaprol and DMPS in the inhibition of procoagulant venom activities, as only moderate inhibition was observed with these metal chelators and no inhibition was found at all for DMPS on *D. russelii* venom. It is, however, possible that the presence of other metal ions in our bioassay (i.e. calcium) may be partly responsible for these reduced inhibitory activities compared with marimastat. Neither DMPS nor dimercaprol inhibited the non-SVMP stimulated anticoagulant venom activities observed across the venoms. Our data further strengthens recent findings suggesting that small molecule inhibitors such as varespladib and marimastat may have broad, cross-species, neutralizing capabilities that make them highly amenable for translation into new ‘generic’ snakebite therapeutics. Given our evidence that both inhibitors have different specificities, our findings further support the concept that a therapeutic combination consisting of both of these Phase II-approved small molecule toxin inhibitors shows great potential as a new broad-spectrum snakebite treatment.

## Supplementary Materials

The supporting information related to this article can be found in “*Supplementary Materials: Neutralizing effects of small molecule inhibitors and metal chelators on coagulopathic Viperinae snake venom toxins*”. The following are available online at www.mdpi.com/xxx/s1, Figure S1: Duplicate bioassay chromatograms of nanofractionated *E. carinatus* venom in the presence of different concentrations of varespladib; Figure S2: Duplicate bioassay chromatograms of nanofractionated *E. carinatus* venom in the presence of different concentrations of marimastat; Figure S3: Duplicate bioassay chromatograms of nanofractionated *E. carinatus* venom in the presence of different concentrations of dimercaprol; Figure S4: Duplicate bioassay chromatograms of nanofractionated *E. carinatus* venom in the presence of different concentrations of DMPS; Figure S5: Duplicate bioassay chromatograms of nanofractionated *E. ocellatus* venom in the presence of different concentrations of varespladib; Figure S6: Duplicate bioassay chromatograms of nanofractionated *E. ocellatus* venom in the presence of different concentrations of marimastat; Figure S7: Duplicate bioassay chromatograms of nanofractionated *E. ocellatus* venom in the presence of different concentrations of dimercaprol; Figure S8: Duplicate bioassay chromatograms of nanofractionated *E. ocellatus* venom in the presence of different concentrations of DMPS; Figure S9: Duplicate bioassay chromatograms of nanofractionated *D. russelii* venom in the presence of different concentrations of varespladib; Figure S10: Duplicate bioassay chromatograms of nanofractionated *D. russelii* venom in the presence of different concentrations of marimastat; Figure S11: Duplicate bioassay chromatograms of nanofarctionated *D. russelii* venom in the presence of different concentrations of dimercaprol; Figure S12: Duplicate bioassay chromatograms of nanofarctionated *D. russelii* venom in the presence of different concentrations of DMPS; Figure S13: Duplicate bioassay chromatograms of nanofractionated *B. arietans* venom in the presence of different concentrations of varespladib; Figure S14: Duplicate bioassay chromatograms of nanofractionated *B. arietans* venom in the presence of different concentrations of marimastat.

## References

1. Gutiérrez, J.M.; Calvete, J.J.; Habib, A.G.; Harrison, R.A.; Williams, D.J.; Warrell, D.A. Snakebite envenoming. Nature Reviews Disease Primers 2017, 3, 1–21.

2. Rogalski, A.; Soerensen, C.; Op den Brouw, B.; Lister, C.; Dashevsky, D.; Arbuckle, K.; Gloria, A.; Zdenek, C.N.; Casewell, N.R.; Gutiérrez, J.M. Differential procoagulant effects of saw-scaled viper (Serpentes: Viperidae: Echis) snake venoms on human plasma and the narrow taxonomic ranges of antivenom efficacies. Toxicology letters 2017, 280, 159–170.

3. Gutiérrez, J.M.; Theakston, R.D.G.; Warrell, D.A. Confronting the neglected problem of snake bite envenoming: the need for a global partnership. PLoS medicine 2006, 3, e150.

4. Calvete, J.J.; Sanz, L.; Angulo, Y.; Lomonte, B.; Gutiérrez, J.M. Venoms, venomics, antivenomics. FEBS letters 2009, 583, 1736–1743.

5. Lu, Q.; Clemetson, J.; Clemetson, K.J. Snake venoms and hemostasis. Journal of Thrombosis and Haemostasis 2005, 3, 1791–1799.

6. Moura-da-Silva, A.; Butera, D.; Tanjoni, I. Importance of snake venom metalloproteinases in cell biology: effects on platelets, inflammatory and endothelial cells. Current Pharmaceutical Design 2007, 13, 2893–2905.

7. Sant’Ana Malaque, C.M.; Gutiérrez, J.M. Snakebite Envenomation in Central and South America. Critical Care Toxicology 2016, 1–22.

8. Maduwage, K.; Isbister, G.K. Current treatment for venom-induced consumption coagulopathy resulting from snakebite. PLoS neglected tropical diseases 2014, 8, e3220.

9. Ainsworth, S.; Slagboom, J.; Alomran, N.; Pla, D.; Alhamdi, Y.; King, S.I.; Bolton, F.M.; Gutiérrez, J.M.; Vonk, F.J.; Toh, C.-H. The paraspecific neutralisation of snake venom induced coagulopathy by antivenoms. Communications biology 2018, 1, 1–14.

10. Slagboom, J.; Kool, J.; Harrison, R.A.; Casewell, N.R. Haemotoxic snake venoms: their functional activity, impact on snakebite victims and pharmaceutical promise. British journal of haematology 2017, 177, 947–959.

11. Kang, T.S.; Georgieva, D.; Genov, N.; Murakami, M.T.; Sinha, M.; Kumar, R.P.; Kaur, P.; Kumar, S.; Dey, S.; Sharma, S. Enzymatic toxins from snake venom: structural characterization and mechanism of catalysis. The FEBS journal 2011, 278, 4544–4576.

12. Kini, R.M. Excitement ahead: structure, function and mechanism of snake venom phospholipase A2 enzymes. Toxicon 2003, 42, 827–840.

13. Matsui, T.; Fujimura, Y.; Titani, K. Snake venom proteases affecting hemostasis and thrombosis. Biochimica et biophysica acta 2000, 1477, 146–156.

14. Serrano, S.M.; Maroun, R.C. Snake venom serine proteinases: sequence homology vs. substrate specificity, a paradox to be solved. Toxicon 2005, 45, 1115–1132.

15. Alvarez-Flores, M.; Faria, F.; de Andrade, S.; Chudzinski-Tavassi, A. Snake venom components affecting the coagulation system. Snake Venoms 2017, 417–436.

16. Ramos, O.; Selistre-de-Araujo, H. Snake venom metalloproteases—structure and function of catalytic and disintegrin domains. Comparative Biochemistry and Physiology Part C: Toxicology & Pharmacology 2006, 142, 328–346.

17. Williams, H.F.; Layfield, H.J.; Vallance, T.; Patel, K.; Bicknell, A.B.; Trim, S.A.; Vaiyapuri, S. The urgent need to develop novel strategies for the diagnosis and treatment of snakebites. Toxins 2019, 11, 363.

18. Bulfone, T.C.; Samuel, S.P.; Bickler, P.E.; Lewin, M.R. Developing small molecule therapeutics for the Initial and adjunctive treatment of snakebite. Journal of tropical medicine 2018, 1–10.

19. Resiere, D.; Gutiérrez, J.M.; Névière, R.; Cabié, A.; Hossein, M.; Kallel, H. Antibiotic therapy for snakebite envenoming. Journal of Venomous Animals and Toxins including Tropical Diseases 2020, 26, 1–2.

20. Albulescu, L.-O.; Hale, M.S.; Ainsworth, S.; Alsolaiss, J.; Crittenden, E.; Calvete, J.J.; Evans, C.; Wilkinson, M.C.; Harrison, R.A.; Kool, J. Preclinical validation of a repurposed metal chelator as an early-intervention therapeutic for hemotoxic snakebite. Science Translational Medicine 2020, 12, eaay8314.

21. Abraham, E.; Naum, C.; Bandi, V.; Gervich, D.; Lowry, S.F.; Wunderink, R.; Schein, R.M.; Macias, W.; Skerjanec, S.; Dmitrienko, A. Efficacy and safety of LY315920Na/S-5920, a selective inhibitor of 14-kDa group IIA secretory phospholipase A2, in patients with suspected sepsis and organ failure. Critical care medicine 2003, 31, 718–728.

22. Nicholls, S.J.; Kastelein, J.J.; Schwartz, G.G.; Bash, D.; Rosenson, R.S.; Cavender, M.A.; Brennan, D.M.; Koenig, W.; Jukema, J.W.; Nambi, V. Varespladib and cardiovascular events in patients with an acute coronary syndrome: the VISTA-16 randomized clinical trial. Jama 2014, 311, 252–262.

23. Lewin, M.; Samuel, S.; Merkel, J.; Bickler, P. Varespladib (LY315920) appears to be a potent, broad-spectrum, inhibitor of snake venom phospholipase A2 and a possible pre-referral treatment for envenomation. Toxins 2016, 8, 248–263.

24. Gutiérrez, J.M.; Lewin, M.R.; Williams, D.; Lomonte, B. Varespladib (LY315920) and Methyl Varespladib (LY333013) Abrogate or Delay Lethality Induced by Presynaptically Acting Neurotoxic Snake Venoms. Toxins 2020, 12, 131.

25. Bryan-Quirós, W.; Fernández, J.; Gutiérrez, J.M.; Lewin, M.R.; Lomonte, B. Neutralizing properties of LY315920 toward snake venom group I and II myotoxic phospholipases A2. Toxicon 2019, 157, 1–7.

26. Bittenbinder, M.A.; Zdenek, C.N.; Op den Brouw, B.; Youngman, N.J.; Dobson, J.S.; Naude, A.; Vonk, F.J.; Fry, B.G. Coagulotoxic cobras: Clinical implications of strong anticoagulant actions of African spitting Naja venoms that are not neutralised by antivenom but are by LY315920 (Varespladib). Toxins 2018, 10, 516–526.

27. Underwood, C.; Min, D.; Lyons, J.; Hambley, T. The interaction of metal ions and Marimastat with matrix metalloproteinase 9. Journal of inorganic biochemistry 2003, 95, 165–170.

28. Peterson, M.; Porter, K.; Loftus, I.; Thompson, M.; London, N. Marimastat Inhibits Neointimal Thickening in aModel of Human Arterial Intimal Hyperplasia. European Journal of Vascular and Endovascular Surgery 2000, 19, 461–467.

29. Curran, S.; Murray, G.I. Matrix metalloproteinases in tumour invasion and metastasis. The Journal of pathology 1999, 189, 300–308.

30. Rasmussen, H.S.; McCann, P.P. Matrix metalloproteinase inhibition as a novel anticancer strategy: a review with special focus on batimastat and marimastat. Pharmacology & therapeutics 1997, 75, 69–75.

31. Evans, J.; Stark, A.; Johnson, C.; Daniel, F.; Carmichael, J.; Buckels, J.; Imrie, C.; Brown, P.; Neoptolemos, J. A phase II trial of marimastat in advanced pancreatic cancer. British journal of cancer 2001, 85, 1865.

32. Winer, A.; Adams, S.; Mignatti, P. Matrix metalloproteinase inhibitors in cancer therapy: turning past failures into future successes. Molecular cancer therapeutics 2018, 17, 1147–1155.

33. Rosenbaum, E.; Zahurak, M.; Sinibaldi, V.; Carducci, M.A.; Pili, R.; Laufer, M.; DeWeese, T.L.; Eisenberger, M.A. Marimastat in the treatment of patients with biochemically relapsed prostate cancer: a prospective randomized, double-blind, phase I/II trial. Clinical cancer research 2005, 11, 4437–4443.

34. King, J.; Zhao, J.; Clingan, P.; Morris, D. Randomised double blind placebo control study of adjuvant treatment with the metalloproteinase inhibitor, Marimastat in patients with inoperable colorectal hepatic metastases: significant survival advantage in patients with musculoskeletal side-effects. Anticancer research 2003, 23, 639–645.

35. Levin, V.A.; Phuphanich, S.; Yung, W.A.; Forsyth, P.A.; Del Maestro, R.; Perry, J.R.; Fuller, G.N.; Baillet, M. Randomized, double-blind, placebo-controlled trial of marimastat in glioblastoma multiforme patients following surgery and irradiation?. Journal of neuro-oncology 2006, 78, 295–302.

36. Bramhall, S.; Schulz, J.; Nemunaitis, J.; Brown, P.; Baillet, M.; Buckels, J. A double-blind placebo-controlled, randomised study comparing gemcitabine and marimastat with gemcitabine and placebo as first line therapy in patients with advanced pancreatic cancer. British journal of cancer 2002, 87, 161.

37. Howes, J.-M.; Theakston, R.D.G.; Laing, G. Neutralization of the haemorrhagic activities of viperine snake venoms and venom metalloproteinases using synthetic peptide inhibitors and chelators. Toxicon 2007, 49, 734–739.

38. Zhang, D.; Botos, I.; Gomis-Rüth, F.-X.; Doll, R.; Blood, C.; Njoroge, F.G.; Fox, J.W.; Bode, W.; Meyer, E.F. Structural interaction of natural and synthetic inhibitors with the venom metalloproteinase, atrolysin C (form d). Proceedings of the National Academy of Sciences 1994, 91, 8447–8451.

39. Nagase, H.; Woessner, J.F. Matrix metalloproteinases. Journal of Biological chemistry 1999, 274, 21491–21494.

40. Rucavado, A.; Escalante, T.; Gutiérrez, J.M.a. Effect of the metalloproteinase inhibitor batimastat in the systemic toxicity induced by Bothrops asper snake venom: understanding the role of metalloproteinases in envenomation. Toxicon 2004, 43, 417–424.

41. Arias, A.S.; Rucavado, A.; Gutiérrez, J.M. Peptidomimetic hydroxamate metalloproteinase inhibitors abrogate local and systemic toxicity induced by Echis ocellatus (saw-scaled) snake venom. Toxicon 2017, 132, 40–49.

42. Layfield, H.J.; Williams, H.F.; Ravishankar, D.; Mehmi, A.; Sonavane, M.; Salim, A.; Vaiyapuri, R.; Lakshminarayanan, K.; Vallance, T.M.; Bicknell, A.B. Repurposing Cancer Drugs Batimastat and Marimastat to Inhibit the Activity of a Group I Metalloprotease from the Venom of the Western Diamondback Rattlesnake, Crotalus atrox. Toxins 2020, 12, 309.

43. Organization, W.H. WHO model list of essential medicines, 20^th^ list (March 2017, amended August 2017). 2017.

44. Tian, R.; Shi, R. Dimercaprol is An Acrolein Scavenger that Mitigates Acrolein-mediated PC-12 Cells Toxicity and Reduces Acrolein in Rat Following Spinal Cord Injury. Journal of Neurochemistry 2017, 141, 708–720.

45. Verma, S.; Kumar, R.; Khadwal, A.; Singhi, S. Accidental inorganic mercury chloride poisoning in a 2-year old child. Indian Journal of Pediatrics 2010, 77, 1153–1155.

46. Kathirgamanathan, K.; Angaran, P.; Lazo-Langner, A.; Gula, L.J. Cardiac conduction block at multiple levels caused by arsenic trioxide therapy. Canadian Journal of Cardiology 2013, 29, 130.e135-130.e136.

47. Yajima, Y.; Kawaguchi, M.; Yoshikawa, M.; Okubo, M.; Tsukagoshi, E.; Sato, K.; Katakura, A. The effects of 2,3-dimercapto-1-propanesulfonic acid (DMPS) and meso-2,3-dimercaptosuccinic acid (DMSA) on the nephrotoxicity in the mouse during repeated cisplatin (CDDP) treatments. Journal of Pharmacological Sciences 2017, 134, 108–115.

48. Aldhaheri, S.R.; Jeelani, R.; Kohan-Ghadr, H.R.; Khan, S.N.; Mikhael, S.; Washington, C.; Morris, R.T.; Abu-Soud, H.M. Dimercapto-1-propanesulfonic acid (DMPS) induces metaphase II mouse oocyte deterioration. Free Radic Biol Med 2017, 112, 445–451.

49. Still, K.; Nandlal, R.S.; Slagboom, J.; Somsen, G.W.; Casewell, N.R.; Kool, J. Multipurpose HTS coagulation analysis: assay development and assessment of coagulopathic snake venoms. Toxins 2017, 9, 382.

50. Xie, C.; Slagboom, J.; Albulescu, L.-O.; Bruyneel, B.; Still, K.; Vonk, F.J.; Somsen, G.W.; Casewell, N.R.; Kool, J. Antivenom Neutralization of Coagulopathic Snake Venom Toxins Assessed by Bioactivity Profiling Using Nanofractionation Analytics. Toxins 2020, 12, 53.

51. Slagboom, J.; Mladić, M.; Xie, C.; Kazandjian, T.D.; Vonk, F.; Somsen, G.W.; Casewell, N.R.; Kool, J. High throughput screening and identification of coagulopathic snake venom proteins and peptides using nanofractionation and proteomics approaches. PLoS Neglected Tropical Diseases 2020, 14, e0007802.

52. Sharma, M.; Das, D.; Iyer, J.K.; Kini, R.M.; Doley, R. Unveiling the complexities of Daboia russelii venom, a medically important snake of India, by tandem mass spectrometry. Toxicon 2015, 107, 266–281.

53. Hiremath, V.; Urs, A.N.; Joshi, V.; Suvilesh, K.; Savitha, M.; Amog, P.U.; Rudresha, G.; Yariswamy, M.; Vishwanath, B. Differential action of medically important Indian BIG FOUR snake venoms on rodent blood coagulation. Toxicon 2016, 110, 19–26.

54. Hiremath, V.; Yariswamy, M.; Nanjaraj Urs, A.; Joshi, V.; Suvilesh, K.; Ramakrishnan, C.; Nataraju, A.; Vishwanath, B. Differential action of Indian BIG FOUR snake venom toxins on blood coagulation. Toxin Reviews 2014, 33, 23–32.

55. Albulescu, L.-O.; Xie, C.; Ainsworth, S.; Alsolaiss, J.; Crittenden, E.; Dawson, C.A.; Softley, R.; Bartlett, K.E.; Harrison, R.A.; Kool, J., et al. A combination of two small molecule toxin inhibitors provides pancontinental preclinical efficacy against viper snakebite. bioRxiv 2020.

56. Wang, Y.; Zhang, J.; Zhang, D.; Xiao, H.; Xiong, S.; Huang, C. Exploration of the inhibitory potential of varespladib for snakebite envenomation. Molecules 2018, 23, 391–403.

57. Kini, R.M.; Sidhu, S.S.; Laustsen, A.H. Biosynthetic oligoclonal antivenom (BOA) for snakebite and next-generation treatments for snakebite victims. Toxins 2018, 10, 534.

58. Knudsen, C.; Ledsgaard, L.; Dehli, R.I.; Ahmadi, S.; Sørensen, C.V.; Laustsen, A.H. Engineering and design considerations for next-generation snakebite antivenoms. Toxicon 2019, 167, 67–75.

59. Lewin, M.R.; Gutiérrez, J.M.; Samuel, S.P.; Herrera, M.; Bryan-Quirós, W.; Lomonte, B.; Bickler, P.E.; Bulfone, T.C.; Williams, D.J. Delayed oral LY333013 rescues mice from highly neurotoxic, lethal doses of Papuan Taipan (Oxyuranus scutellatus) venom. Toxins 2018, 10, 380–386.

60. Lewin, M.R.; Gilliam, L.L.; Gilliam, J.; Samuel, S.P.; Bulfone, T.C.; Bickler, P.E.; Gutiérrez, J.M. Delayed LY333013 (oral) and LY315920 (intravenous) reverse severe neurotoxicity and rescue juvenile pigs from lethal doses of Micrurus fulvius (Eastern Coral snake) venom. Toxins 2018, 10, 479.

61. Harrison, R.A.; Oluoch, G.O.; Ainsworth, S.; Alsolaiss, J.; Bolton, F.; Arias, A.-S.; Gutiérrez, J.-M.; Rowley, P.; Kalya, S.; Ozwara, H. Preclinical antivenom-efficacy testing reveals potentially disturbing deficiencies of snakebite treatment capability in East Africa. PLoS neglected tropical diseases 2017, 11, e0005969–e0005969.

62. Sharma, S.K.; Chappuis, F.; Jha, N.; Bovier, P.A.; Loutan, L.; Koirala, S. Impact of snake bites and determinants of fatal outcomes in southeastern Nepal. The American journal of tropical medicine and hygiene 2004, 71, 234–238.

63. Abubakar, S.; Habib, A.; Mathew, J. Amputation and disability following snakebite in Nigeria. Tropical Doctor 2010, 40, 114–116.

