## Supplementary Material for "Neutralizing effects of small molecule inhibitors and metal chelators on coagulopathic *Viperinae* snake venom toxins"

S1 Inhibitory effects of varespladib, marimastat, dimercaprol and DMPS on coagulopathic venom toxins from *Echis* species

The *Echis* species investigated in this study were *Echis carinatus* (India) and *Echis ocellatus* (Nigeria). Venoms from these snakes were analyzed to assess the inhibition of coagulopathic venom toxins by different concentrations of varespladib, marimastat, dimercaprol and DMPS. Duplicate reconstructed coagulation bioassay chromatograms showing the effectiveness of the tested inhibitors at various concentrations against coagulopathic toxins from nanofractionated *E. carinatus* venom are shown in Figures S1-S4. In each Figure, the bioassay chromatograms depicted on the right, and which do not have superimposed correlated UV data, represent replicates of the chromatograms on the left. Each assay was kinetically measured for 100 min. The coagulation curves were plotted versus the chromatographic retention time for each fraction collected in three different ways to fully depict both the procoagulant and anticoagulant activities in each well. The slope of the average 0-5 min reading was used to depict very fast coagulation activity, the slope of the average 0-20 min reading was used for slightly/medium increased coagulation activity, and the single reading at 100 min was used to assess anticoagulation activity. This way, the procoagulant activity is presented in two different distinct bioactivity chromatograms to clearly discriminate between very fast coagulation (i.e. maximum absorbance and thus full coagulation reached within a few minutes) and slightly/medium increased coagulation (i.e. maximum absorbance reached within tens of minutes).

Varespladib inhibited both anticoagulant and procoagulant venom activities (Figure S1). In the venom-only analysis, a weak positive peak (22.1 min) followed by an intense sharp positive peak (22.3 min) was observed for the very fast coagulation activity. Two sets of intense and broad positive peaks (19.9-21.2 min and 21.2-22.8 min), which were the result of several closely co-eluting proteins, were observed for the slightly/medium increased coagulation activity. An intense sharp negative peak (19.4-19.9 min) was observed in the anticoagulation chromatogram. Upon increasing the varespladib concentration, all bioactivity peaks decreased and became narrower. For the very fast coagulation activity, the weak positive peak (22.1 min) was readily neutralized by 0.8 μM varespladib. No activity was detected at a 4 μM varespladib concentration. For the slightly/medium increased coagulation activity, the first cluster of positive peaks (19.9-21.2 min) was fully neutralized at a varespladib concentration of 4 μM, while the activity of the second positive cluster (21.2-22.8 min) decreased in a concentration-dependent manner. Only an intense sharp positive peak (21.9 min) was retained at the highest varespladib concentration tested (20 μM). For anticoagulant activity, the activity of the negative peak (19.4-19.9 min) decreased in both height and width with increasing varespladib concentrations. No activity was detected at 20 μM varespladib.

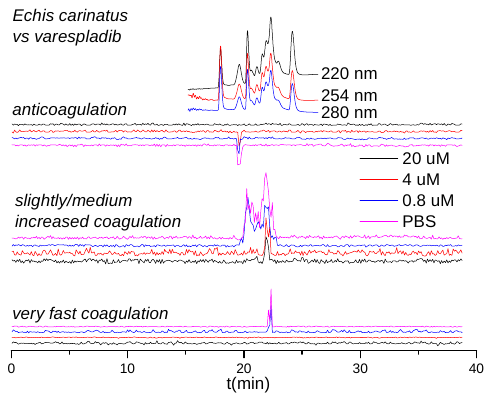

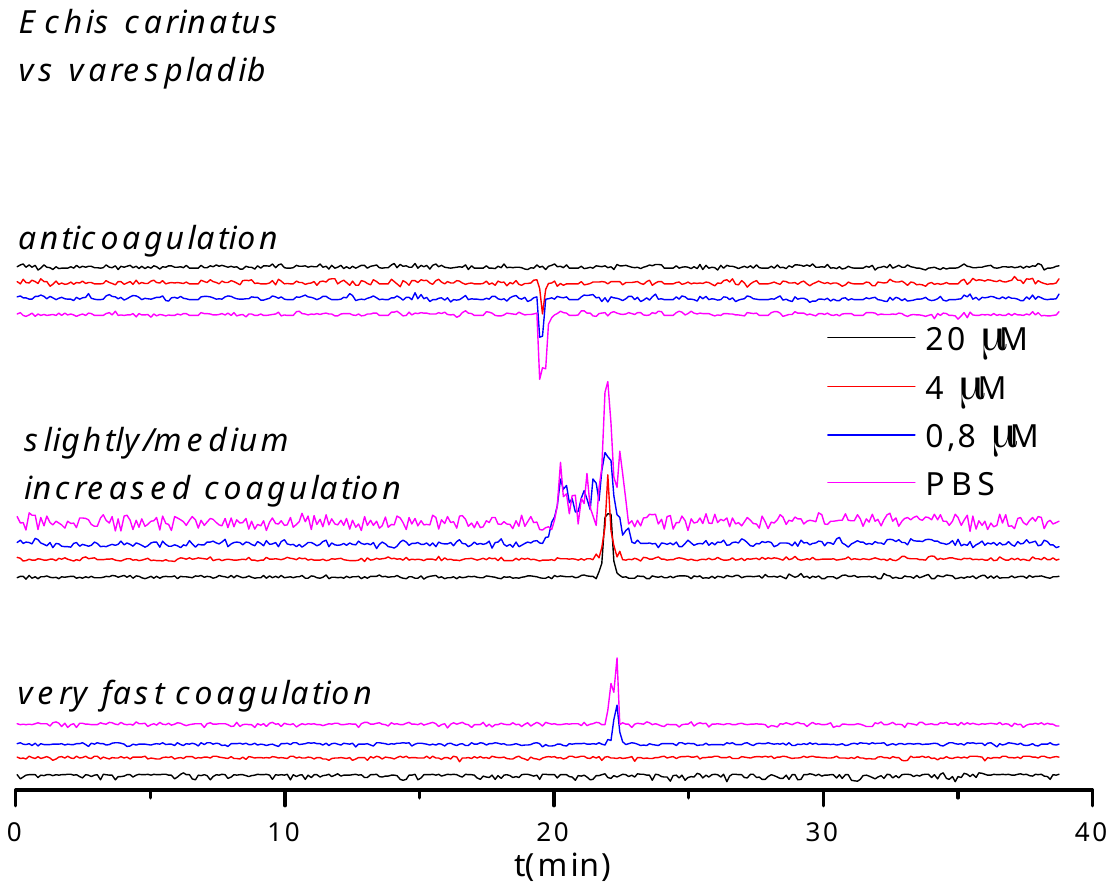

**Figure S1.** Duplicate bioassay chromatograms of nanofractionated *E. carinatus* venom in the presence of different concentrations of varespladib.

Marimastat strongly inhibited procoagulant activity, while displaying limited efficacy against anticoagulant activity, as shown in Figure S2. In the venom-only analysis, a positive shoulder peak indicating potent activity was detected for both the very fast coagulation activity (22.1-22.9 min) and the slightly/medium increased coagulation activity (21.2-23.1 min), and an intense sharp negative peak (19.2-19.9 min) followed by a weak negative peak (20.3 min) was observed in the anticoagulation chromatogram. By increasing the marimastat concentration, the procoagulant activity decreased significantly, while the anticoagulant activity was not affected. For the very fast coagulation activity, only a weak positive peak (22.1-22.9 min) was detected at a marimastat concentration of 0.032 μM, and no activity was observed at a 0.16 μM marimastat concentration. For the slightly/medium increased coagulation activity, the intense positive shoulder peak (21.2-23.1 min) was decreased and became narrower with increasing the marimastat concentrations. No activity was observed at 0.8 μM marimastat. Neither the intense sharp negative peak (19.2-19.9 min) nor the weak negative peak (20.3 min) in the anticoagulation chromatograms were diminished at any marimastat concentration tested.

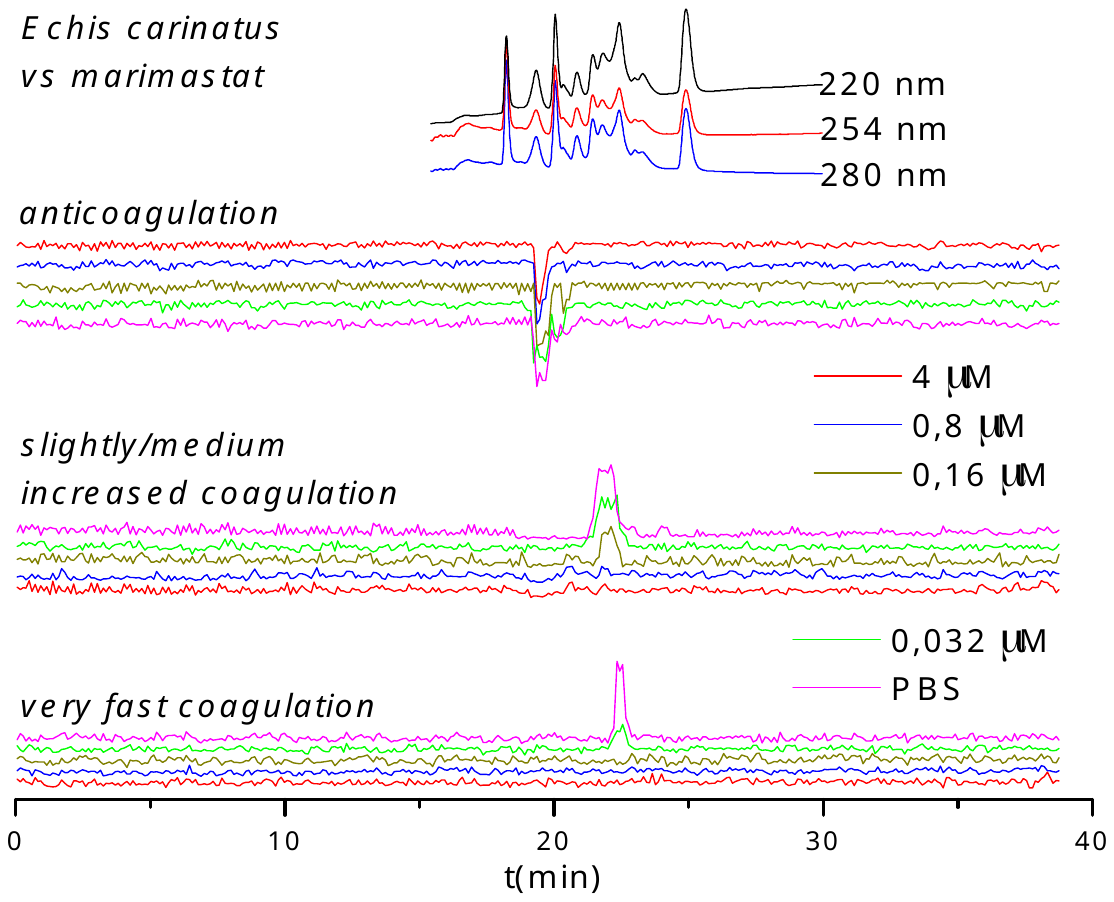

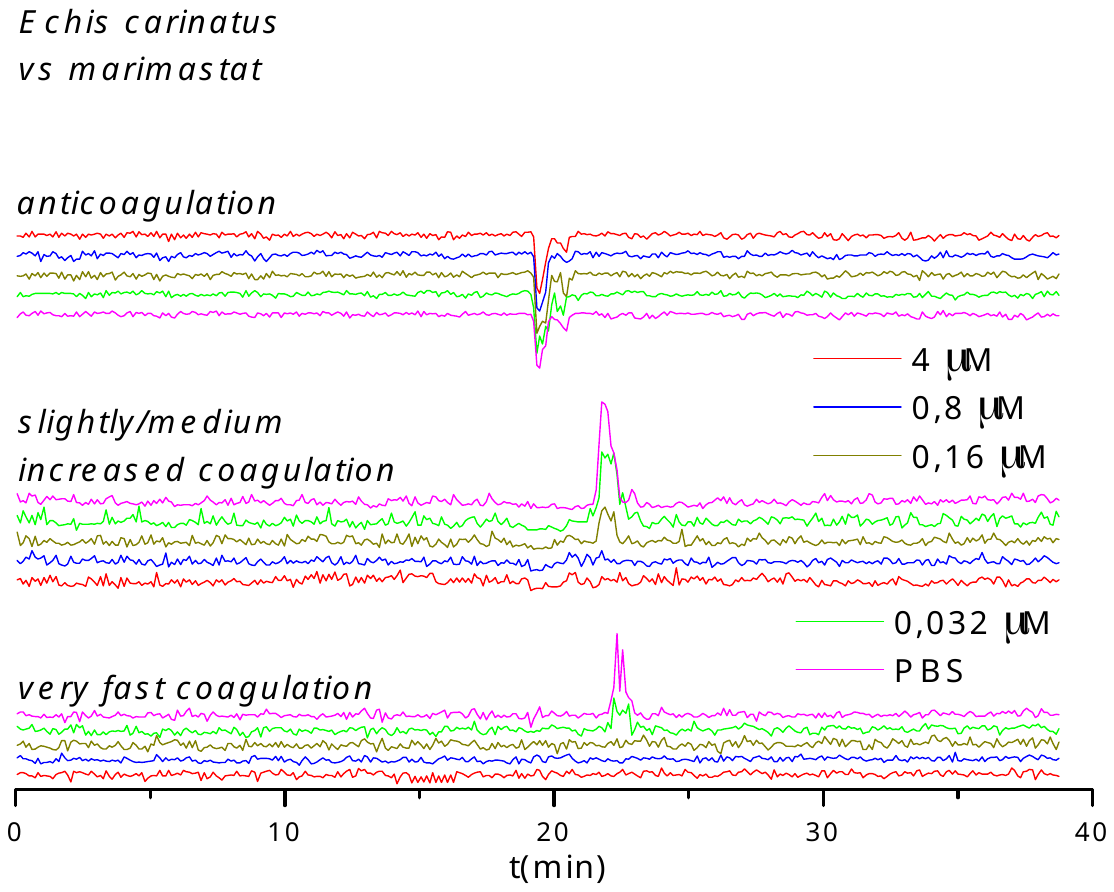

**Figure S2.** Duplicate bioassay chromatograms of nanofractionated *E. carinatus* venom in the presence of different concentrations of marimastat.

The inhibitory effects of dimercaprol on *E. carinatus* venom are shown in Figure S3. In the venom-only analysis, an intense positive shoulder peak (22.0-22.4 min) was noted in the very fast coagulation chromatogram. A weak positive peak (21.5 min) followed by an intense positive peak (21.8 min) and another weak positive peak (22.3 min) made up a non-baseline separated positive peak (21.3-22.4 min), which was followed by a weak peak (22.9 min) in the slightly/medium increased coagulation chromatogram. An intense and relatively broad anticoagulant negative peak (19.1-19.9 min) was also detected. The anticoagulant activity was not influenced by dimercaprol. The activity of the positive shoulder peak (22.0-22.4 min) in the very fast pro-coagulation chromatogram was fully inhibited by 4 μM dimercaprol. For the slightly/medium increased coagulation activity, the majority of non-baseline separated positive peaks (21.3-22.4 min) were neutralized by 0.8 μM dimercaprol, and only a sharp and intense positive peak (22.3 min) was retained at this concentration. No further neutralization was observed upon increasing the concentration of dimercaprol. The latest weak positive peak (22.9 min) which eluted after the non-baseline separated positive peak (21.3-22.4 min) was still visible at a dimercaprol concentration of 0.8 μM, but was fully inhibited by 4 μM dimercaprol.

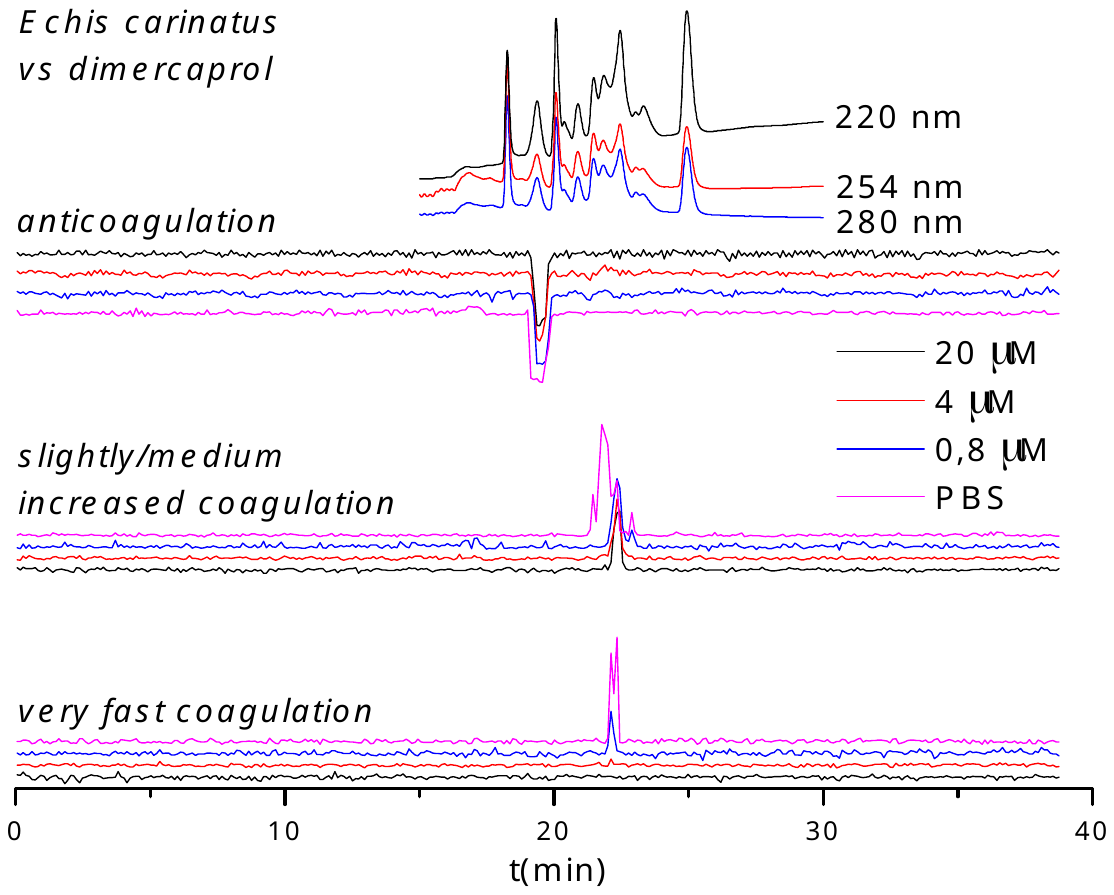

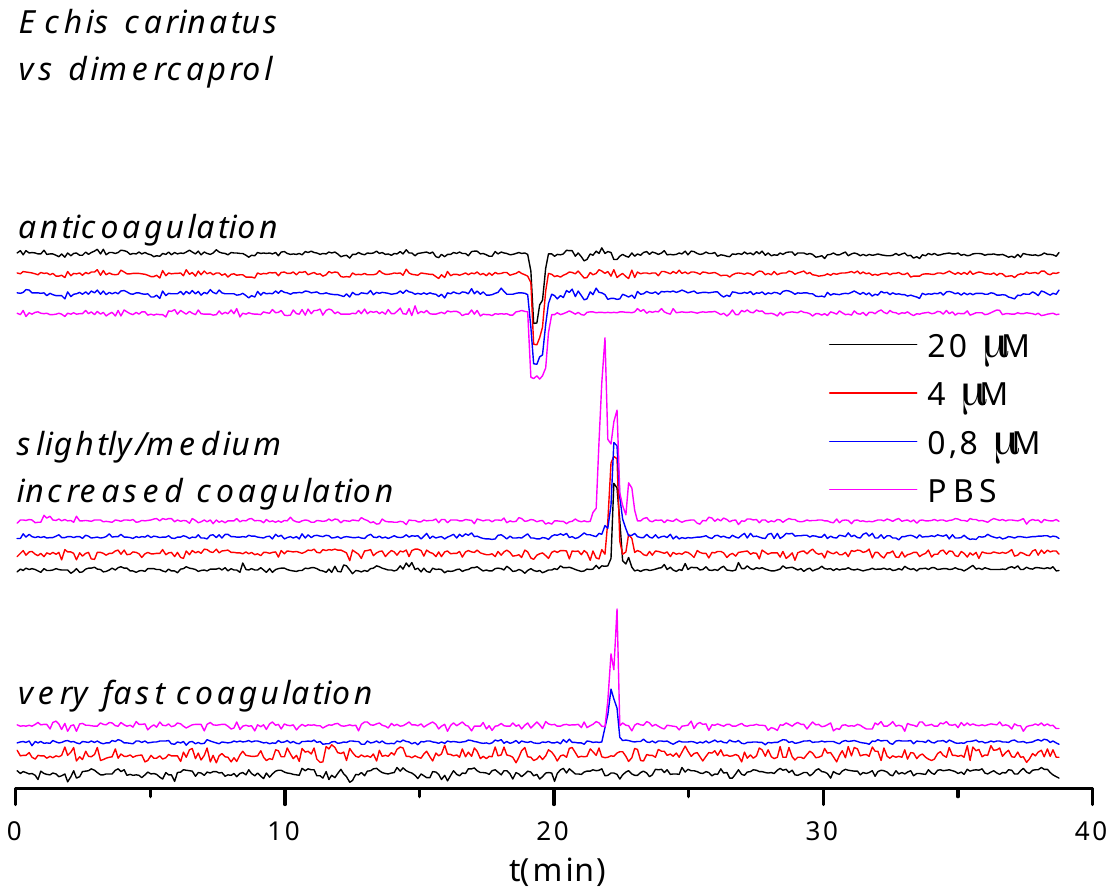

**Figure S3.** Duplicate bioassay chromatograms of nanofractionated *E. carinatus* venom in the presence of different concentrations of dimercaprol.

The inhibitory effect of DMPS on *E. carinatus* venom is shown in Figure S4. Similar to dimercaprol, DMPS showed almost no effect on anticoagulant activity, as expected for these metal-chelating drugs targeting SVMPs. For the very fast coagulation activity, the intense positive shoulder peak (22.0-22.4 min) was reduced to a very weak positive peak (22.1 min) at a concentration of 0.8 μM DMPS. No activity was detected at 4 μM DMPS. For the slightly/medium increased coagulation activity, the non-baseline separated positive peak (21.3-22.4 min) changed into a shoulder peak (21.6-22.7 min) at 0.8 μM DMPS, and was further decreased by increasing the DMPS concentration, despite the fact that full neutralization was not achieved at the highest DMPS concentration tested. The latest eluting weak positive peak (22.9 min) was reduced, but still visible at 0.8 μM DMPS concentration, and was fully inhibited at 4 μM DMPS.

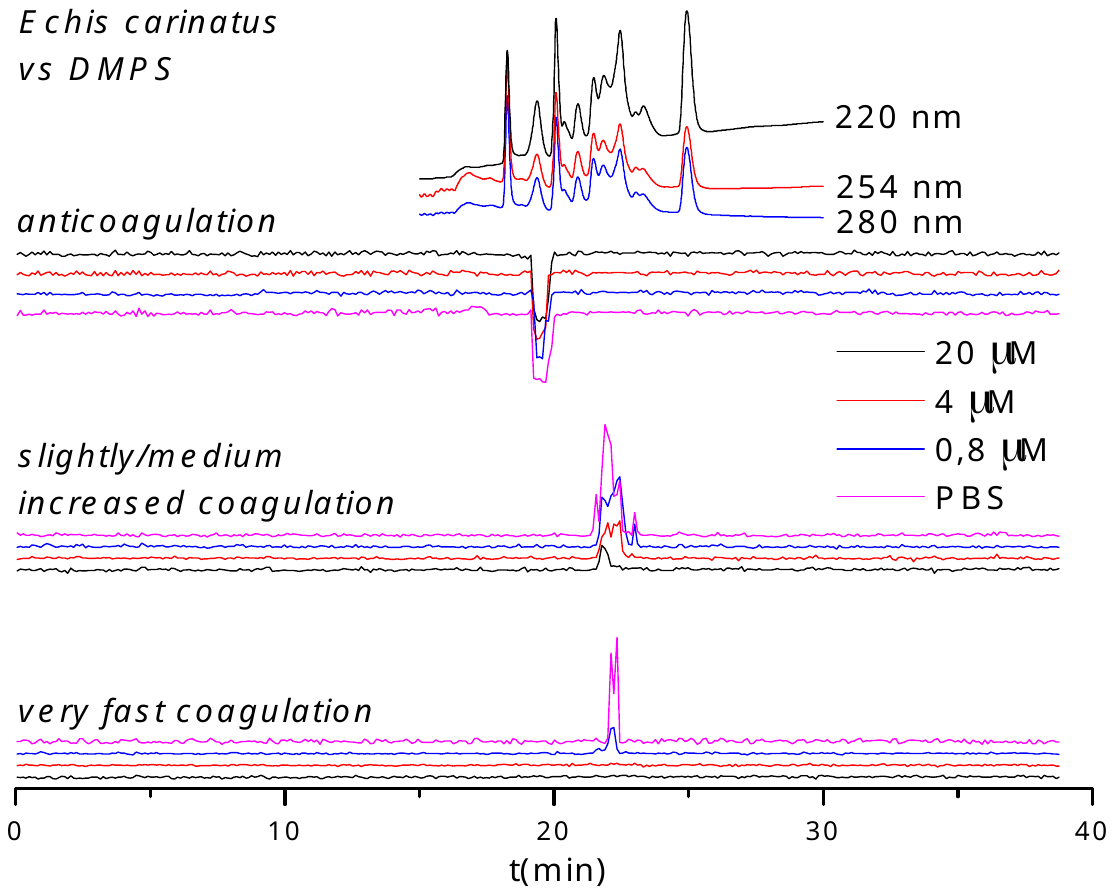

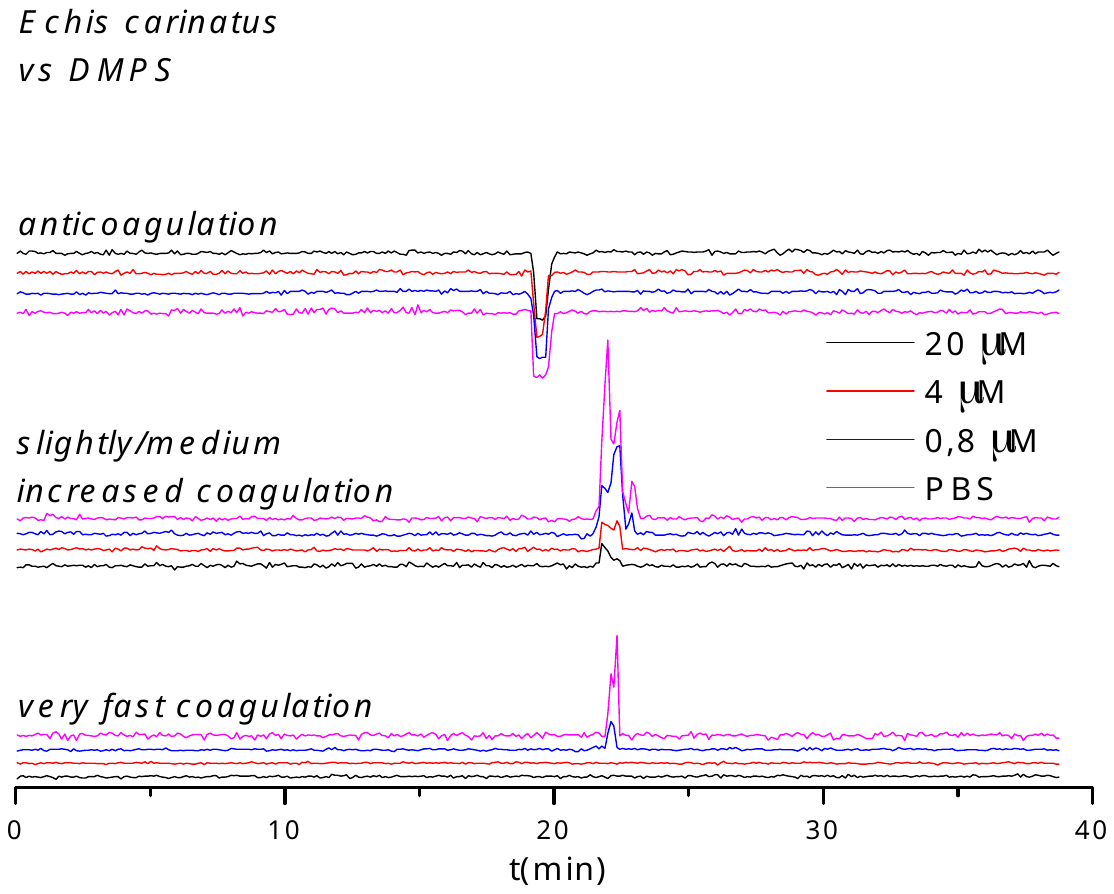

**Figure S4.** Duplicate bioassay chromatograms of nanofractionated *E. carinatus* venom in the presence of different concentrations of DMPS.

The reconstructed coagulation bioassay chromatograms showing the inhibitory effects of different concentrations of varespladib, marimastat, dimercaprol and DMPS against nanofractionated venom toxins from *E. ocellatus* are shown in Figures S5-S8. The inhibitory effect of varespladib on *E.* *ocellatus* venom is shown in Figure S5. For the venom-only analysis, a weak positive peak (24.8 min) followed by an intense positive peak (25.1-26.2 min) was observed in the very fast coagulation chromatogram, an intense positive shoulder peak (25.1-27.1 min) was observed in the slightly/medium increased coagulation chromatogram, and an intense negative peak (23.4-24.4 min) followed by a weak negative peak (26.1 min) was observed in the anticoagulation chromatogram. In presence of varespladib, the weak positive peak (24.8 min) in the very fast coagulation chromatogram was fully neutralized at a low varespladib concentration (0.16 μM). The activity of the intense positive peak (25.1-26.2 min) in the very fast coagulation chromatogram decreased with increasing concentrations of varespladib, but this intense positive peak (52.1-26.2 min) could not be fully neutralized by the highest varespladib concentration tested (20 μM). For the slightly/medium increased coagulation activity, the intense positive shoulder peak (25.1-27.1 min) became smaller and narrower upon increasing the varespladib concentration, and became a sharp intense positive peak (26.0 min) at a concentration of 0.16 μM varespladib. The activity of this intense sharp positive peak (26.0 min) decreased dose-dependently by further increasing the varespladib concentration but full neutralization was not achieved. For the anticoagulant activity, the intense negative peak (23.4-24.4 min) decreased by increasing the varespladib concentration and was fully neutralized by varespladib at a concentration of 4 μM. The activity in the latest eluting weak negative peak (26.1 min) was unaffected by varespladib in the 0.16-4 μM range, but was fully neutralized by 20 μM varespladib.

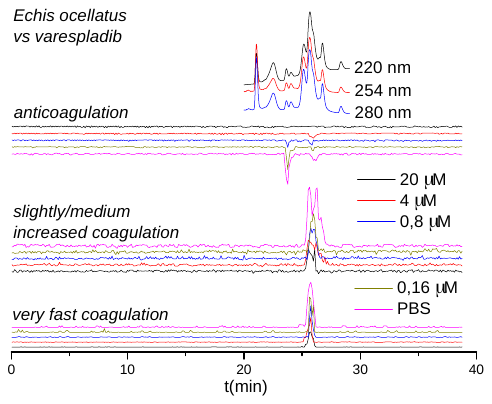

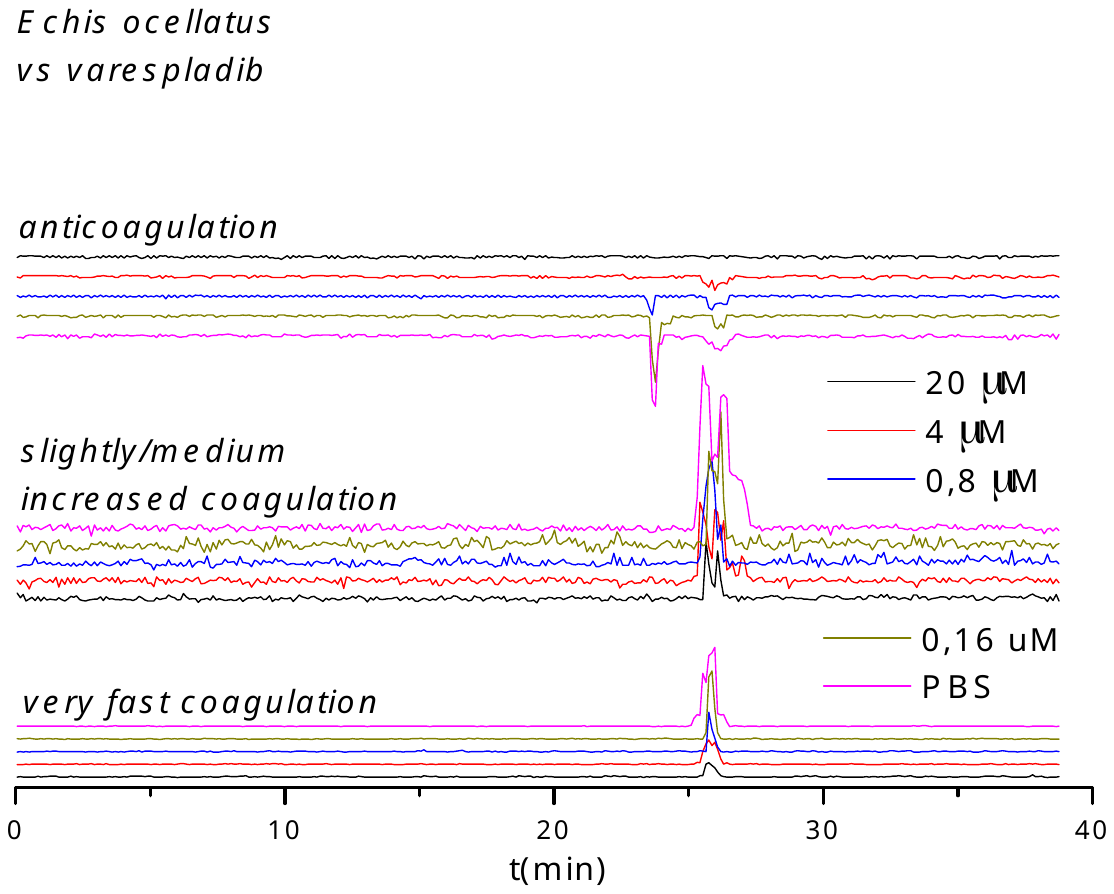

**Figure S5.** Duplicate bioassay chromatograms of nanofractionated *E. ocellatus* venom in the presence of different concentrations of varespladib.

The inhibitory effect of marimastat on *E. ocellatus* venom is shown in Figure S6. In the venom-only analysis, an intense sharp positive peak (25.4-26.2 min) can be seen both in the very fast coagulation activity chromatogram and in the slightly/medium increased coagulation activity chromatogram. An intense sharp negative peak (23.5-24.1 min) can be seen in the anticoagulation activity chromatogram. For the marimastat analyses, the intense sharp negative peak (23.5-24.1 min) in the anticoagulation chrontogram was not influenced by marimastat, while the activity of the intense sharp positive peak (25.4-26.2 min) indicating slightly/medium increased coagulation decreased with increasing marimastat concentrations. Full inactivation of these procoagulant activities was achieved at a marimastat concentration of 4 μM. For the very fast coagulation activity, a weak positive peak (25.8 min) was still observed at 0.032 μM marimastat, which was fully neutralized by 0.16 μM marimastat.

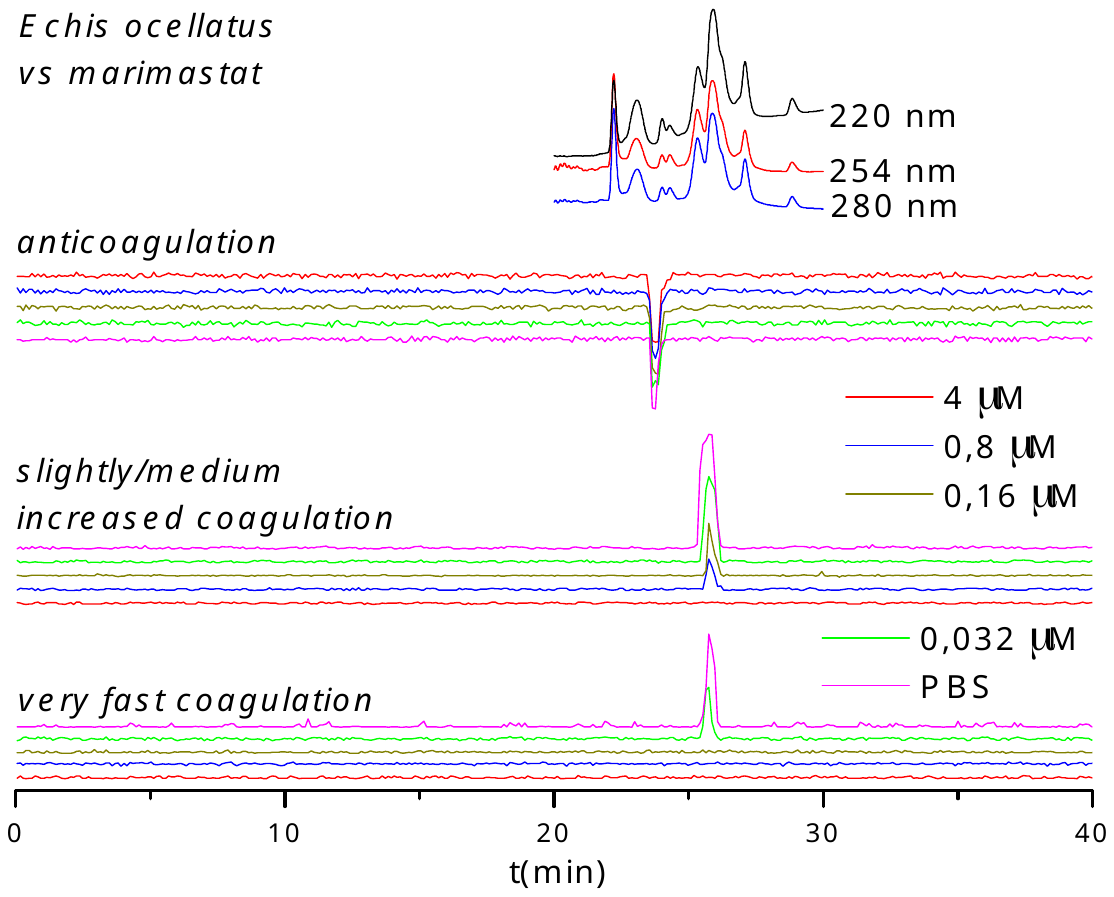

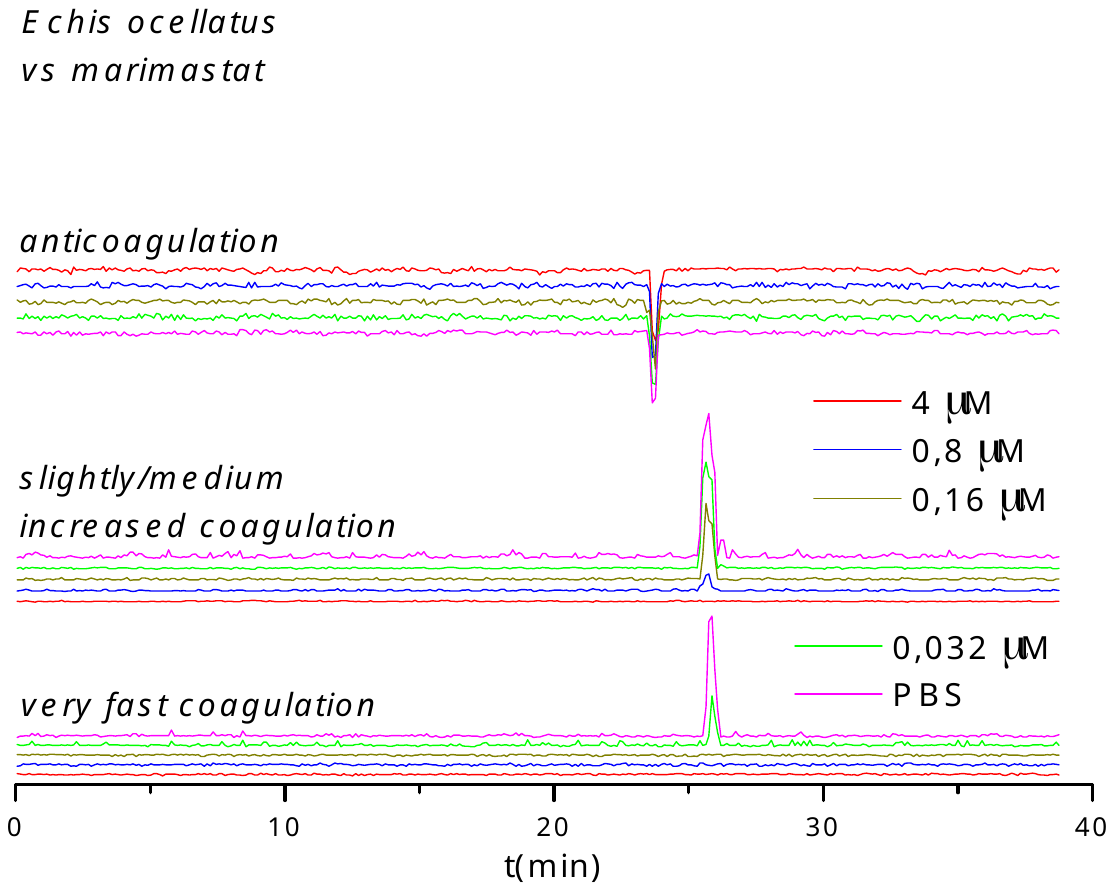

**Figure S6.** Duplicate bioassay chromatograms of nanofractionated *E. ocellatus* venom in the presence of different concentrations of marimastat.

The inhibitory effects of dimercaprol on *E. ocellatus* venom is shown in Figure S7. Only moderate inhibition was achieved against both the very fast coagulation activity (25.2-26.2 min) and the slightly/medium increased coagulation activity (25.2-26.2 min). The positive peaks (25.2-26.2 min) decreased with increasing dimercaprol concentrations, but full inhibition was not achieved. Dimercaprol could not inhibit the anticoagulant activity (23.5-24.1 min).

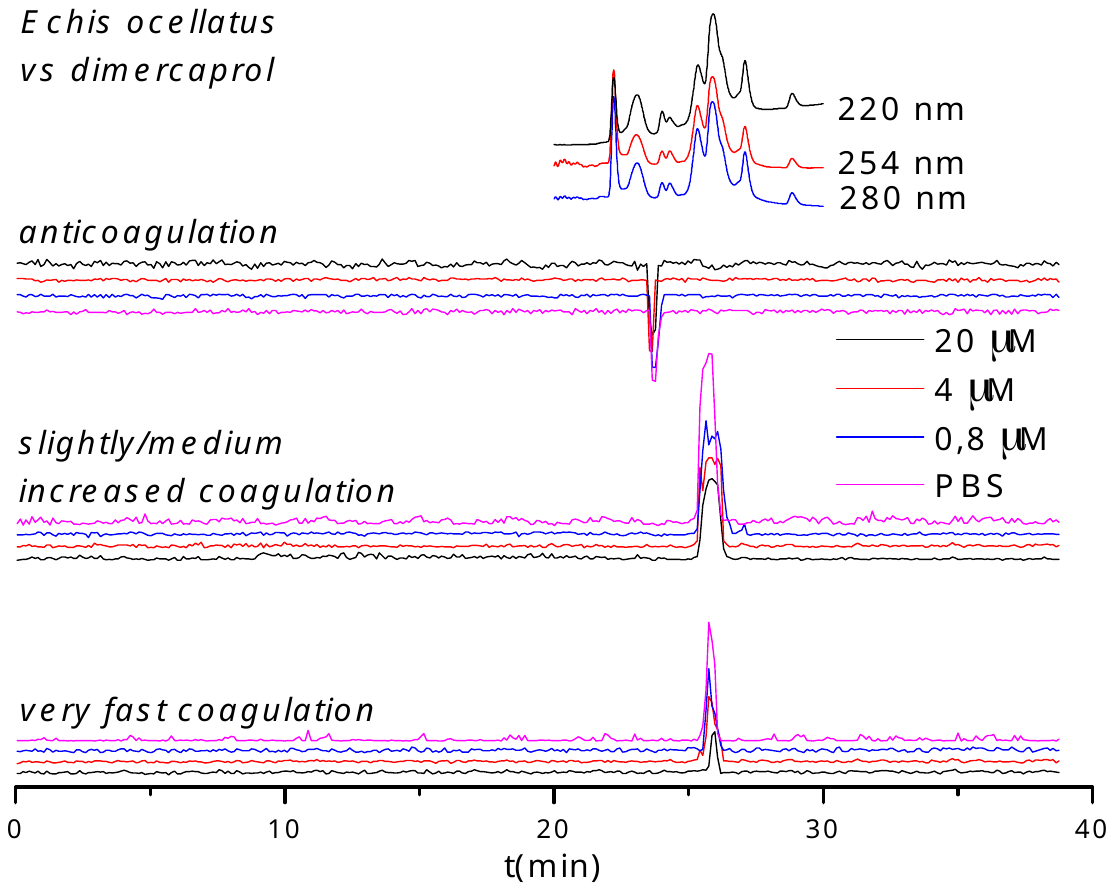

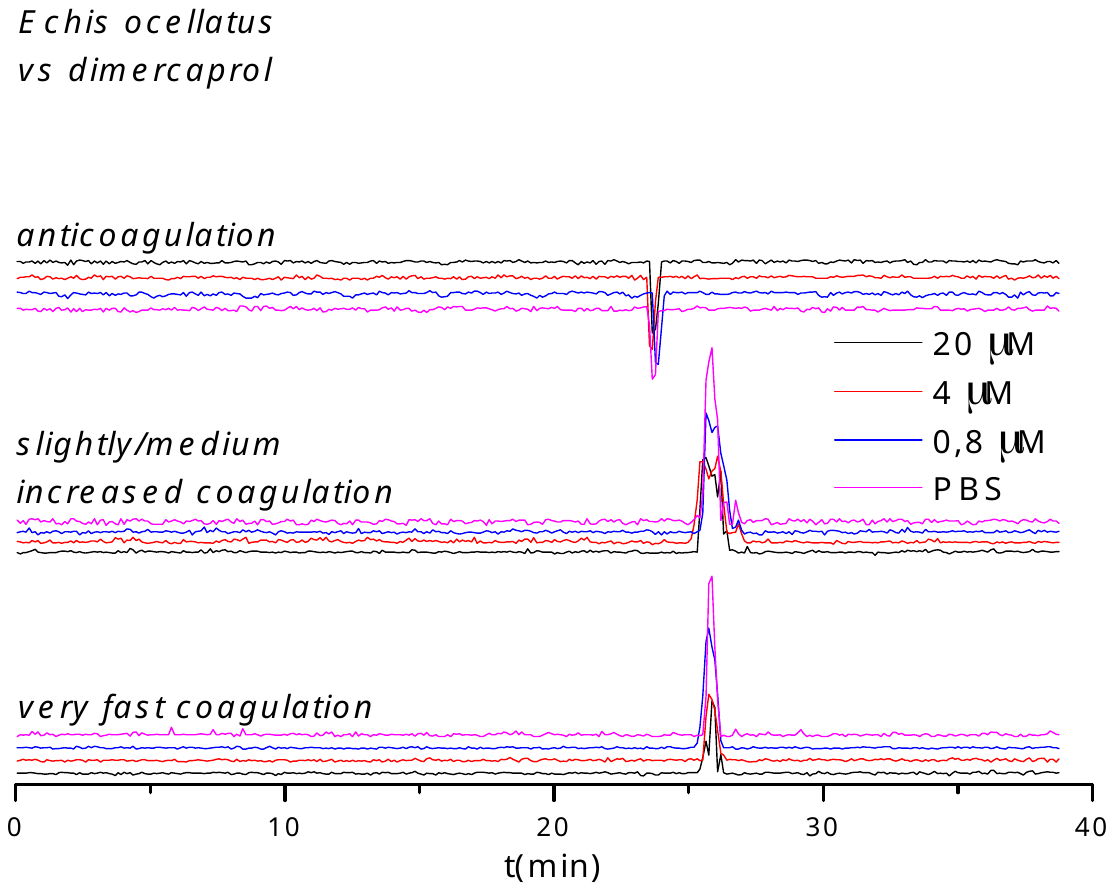

**Figure S7.** Duplicate bioassay chromatograms of nanofractionated *E. ocellatus* venom in the presence of different concentrations of dimercaprol.

The inhibitory effect of DMPS on *E. ocellatus* venom is shown in Figure S8. For the venom-only analysis, an intense sharp positive peak (25.6-26.3 min) was observed both in the very fast coagulation chromatogram and in the slightly/medium increased coagulation chromatogram, and a sharp intense negative peak (23.4-24.1 min) was noted in the anticoagulation chromatogram. The sharp positive peak (25.6-26.3 min) in both the very fast coagulation chromatogram and the slightly/medium increased coagulation chromatogram was reduced by increasing the DMPS concentration. Full inhibition was achieved for the very fast coagulation activity at 20 μM DMPS but was not attained for the slightly/medium increased coagulation activity. The anticoagulant activity (23.4-24.1 min) was not influenced by DMPS.

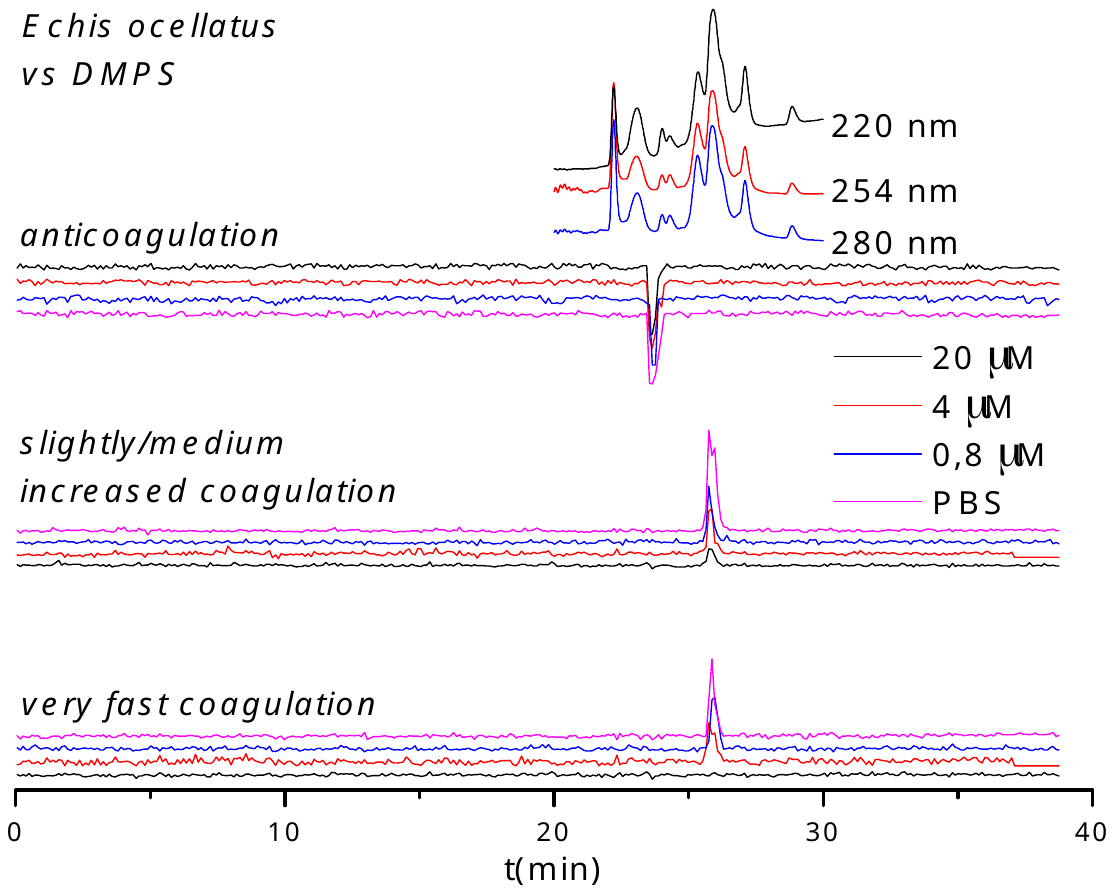

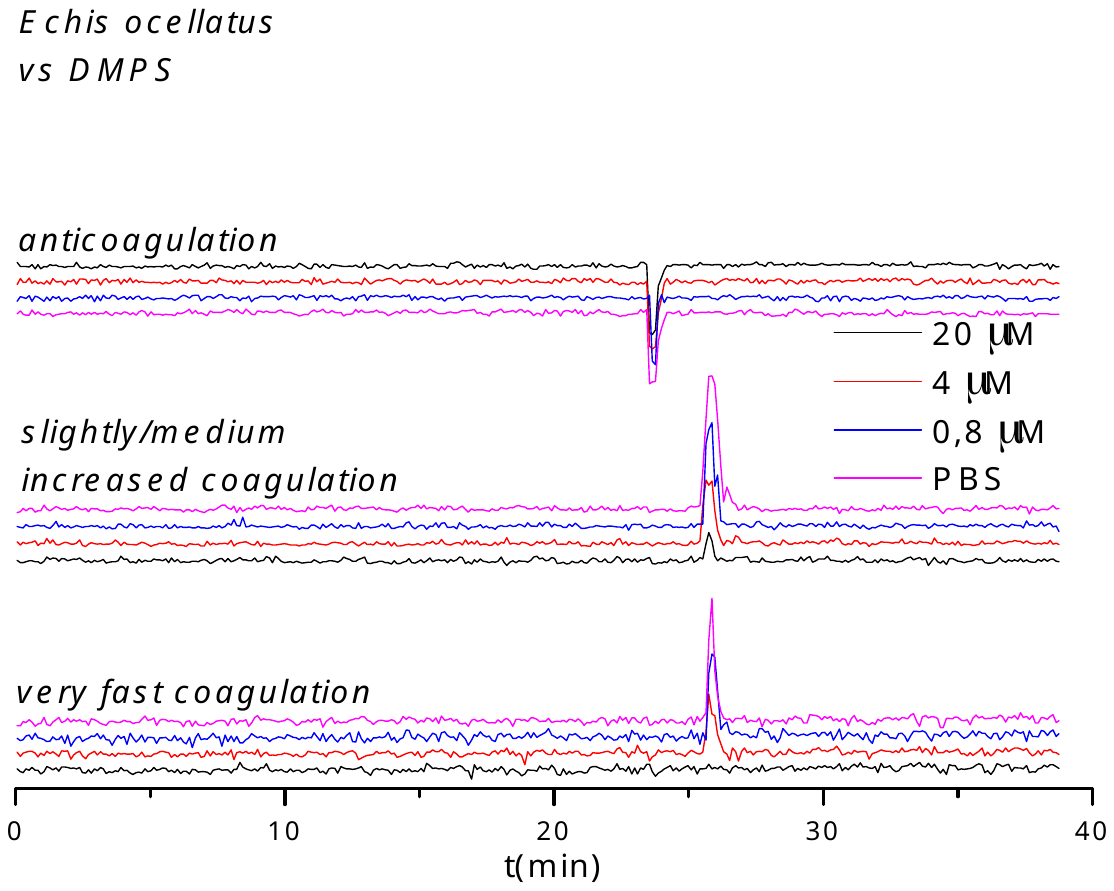

**Figure S8.** Duplicate bioassay chromatograms of nanofractionated *E. ocellatus* venom in the presence of different concentrations of DMPS.

S2 Inhibitory effects of varespladib marimastat, dimercaprol and DMPS on the activities of coagulopathic venom toxins from *Daboia russelii*

The reconstructed duplicate chromatograms showing the inhibitory effects of varespladib, marimastat, dimercaprol and DMPS on nanofractionated *D. russelii* (Sri Lanka) venom are shown in Figures S9-S12. Varespladib could dose-dependently reduce all coagulopathic activities observed, but was not able to fully neutralize these activities at the highest concentration tested (20 μM). Marimastat and dimercaprol could only reduce the pro-coagulopathic activities in a concentration-dependent manner. Marimastat neutralized all pro-coagulant activities at a concentration of 4 μM. Dimercaprol could fully neutralize these activities at the highest concentration tested (20 μM). DMPS showed no inhibition on these activities at tested concentrations of 20 μM and 4 μM.

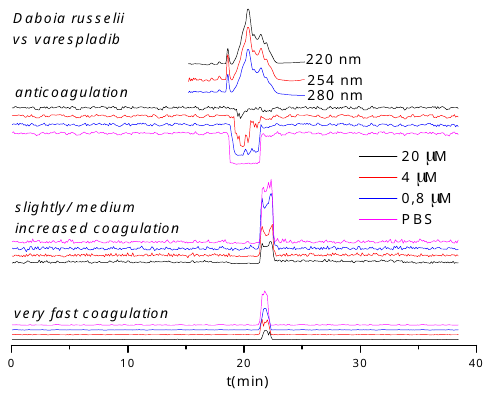

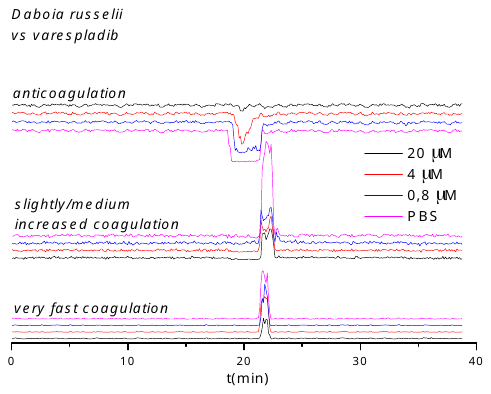

**Figure S9.** Duplicate bioassay chromatograms of nanofractionated *D. russelii* venom in the presence of different concentrations of varespladib.

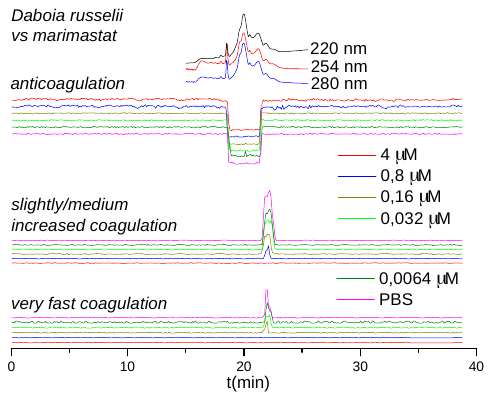

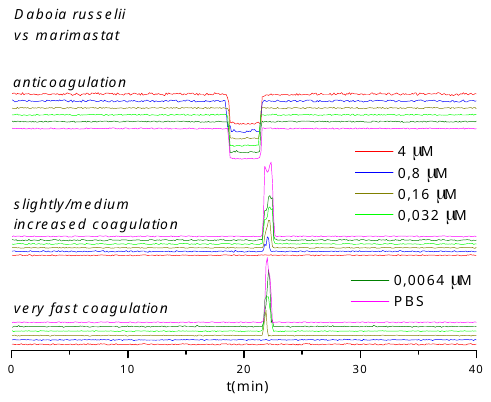

**Figure S10.** Duplicate bioassay chromatograms of nanofractionated *D. russelii* venom in the presence of different concentrations of marimastat.

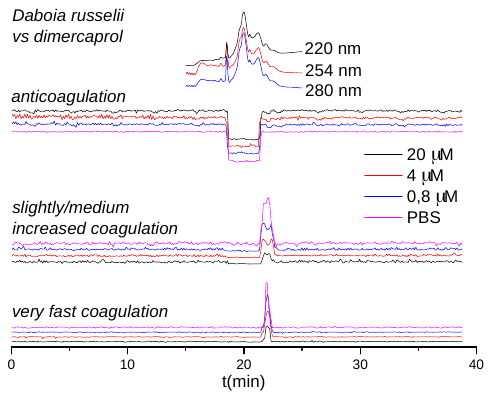

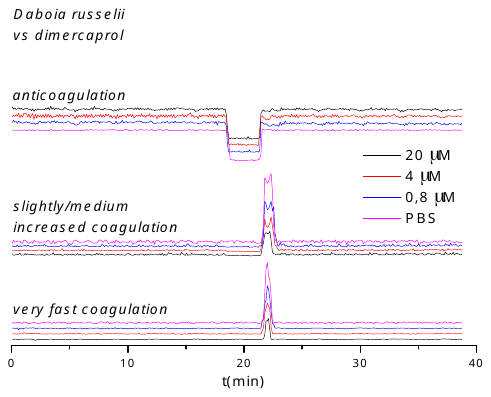

**Figure S11.** Duplicate bioassay chromatograms of nanofarctionated *D. russelii* venom in the presence of different concentrations of dimercaprol.

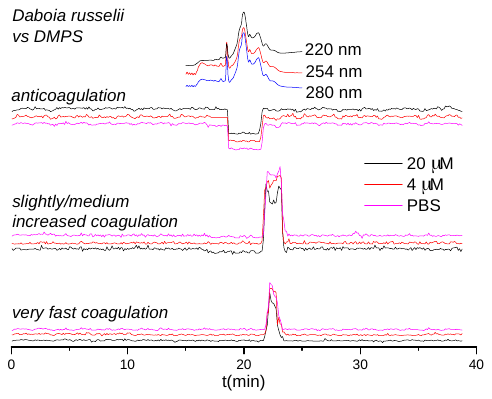

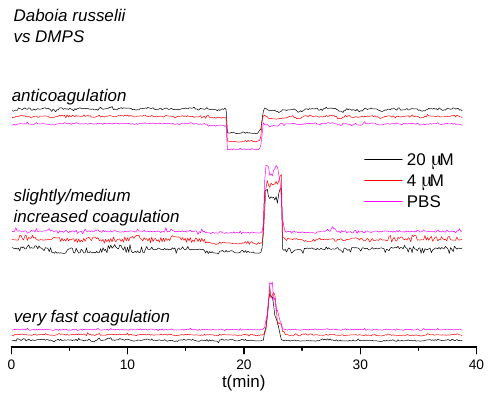

**Figure S12.** Duplicate bioassay chromatograms of nanofarctionated *D. russelii* venom in the presence of different concentrations of DMPS.

S3 Inhibitory effects of varespladib and marimastat on the activities of coagulopathic venom toxins from *Bitis arietans*

The reconstructed duplicate chromatograms showing the inhibitory effects of varespladib and marimastat on nanofractionated *B. arietans* (Nigeria) venom are shown in Figures S13-S14. At the venom concentration used for the analyses (1 mg/ml), only anticoagulant activity was observed. Varespladib could dose-dependently reduce all coagulopathic activities, and full neutralization was achieved at a low concentration (0.8 μM). Marimastat did not affect anticoagulant activity, even at the highest concentration tested (20 μM).

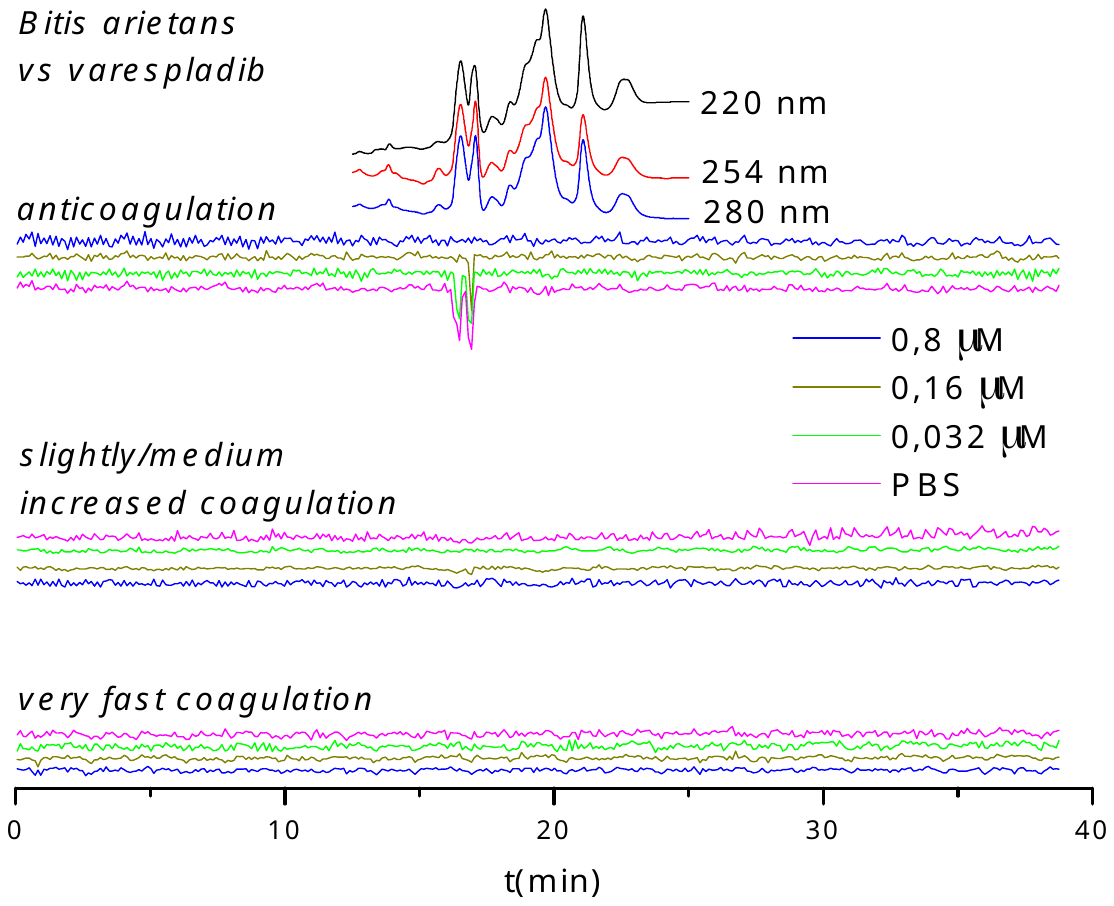

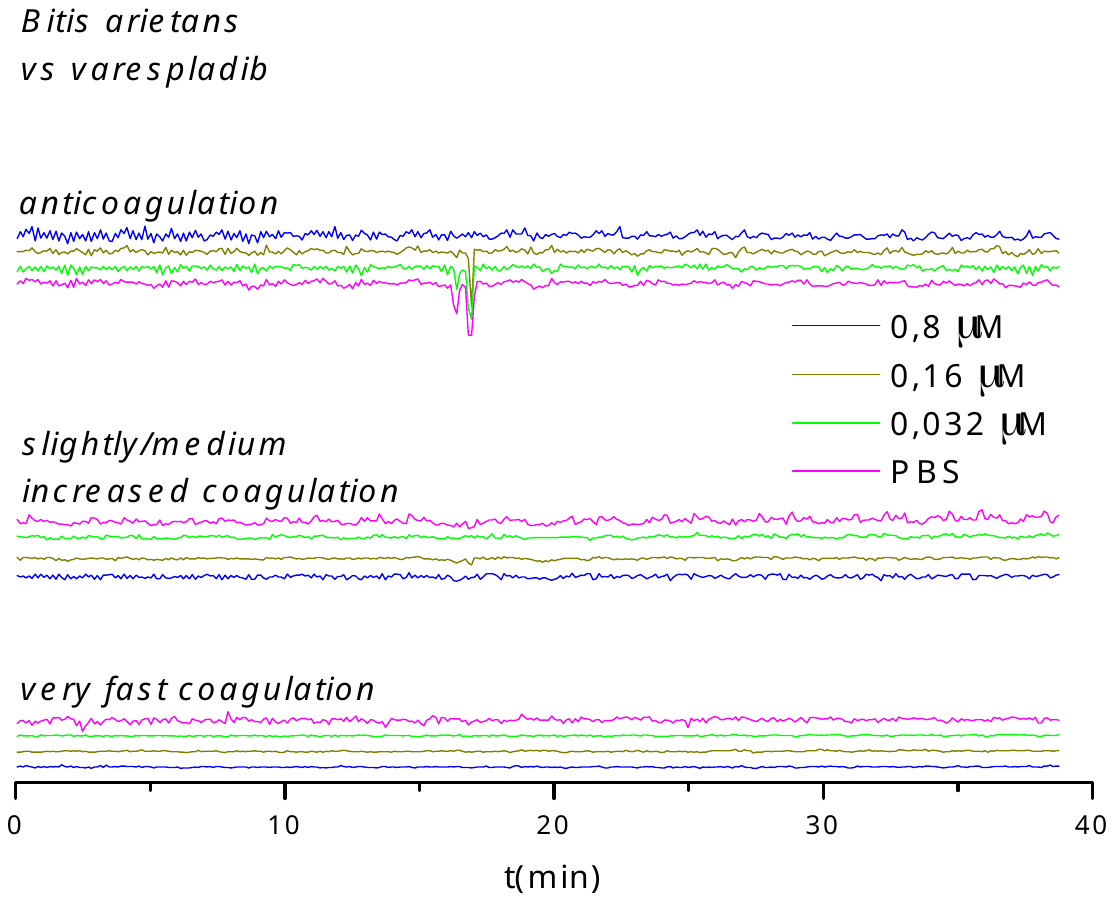

**Figure S13.** Duplicate bioassay chromatograms of nanofractionated *B. arietans* venom in the presence of different concentrations of varespladib.

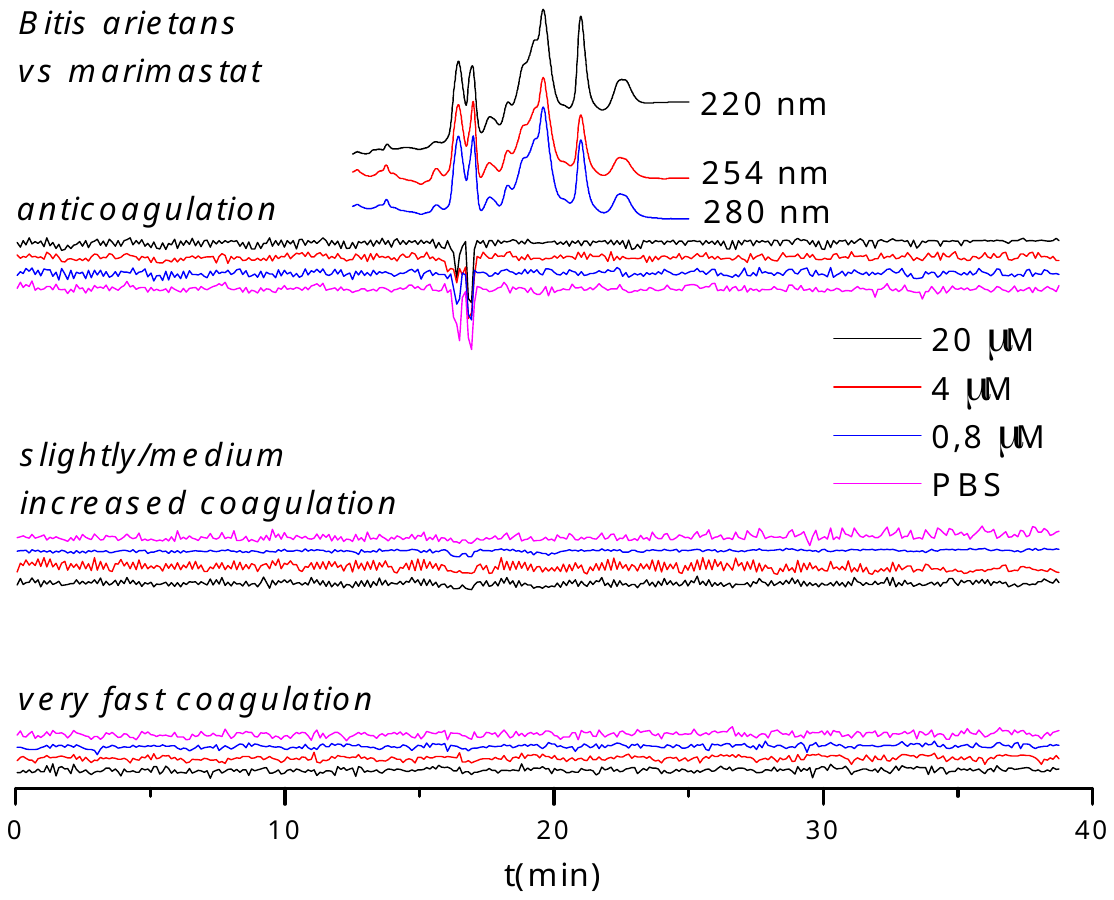

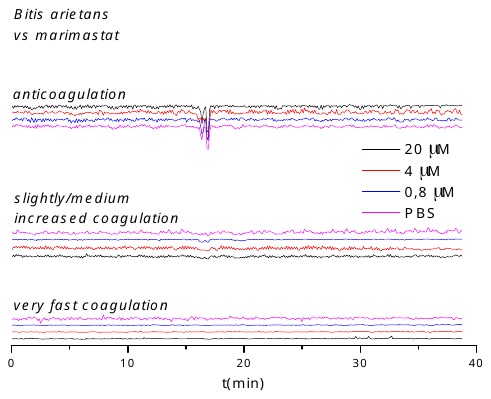

**Figure S14.** Duplicate bioassay chromatograms of nanofractionated *B. arietans* venom in the presence of different concentrations of marimastat.

S4. Inhibitory efficacy of small molecule inhibitors and metal chelators on venom toxins identified and depicted in Table 1.

PA2A1_ECHCA, which was reported to lack hemorrhagic activity [1], was neutralized by 20 μM varespladib, but not affected by marimastat, dimercaprol or DMPS. PA2A5_ECHOC, for which no anticoagulant activity has been reported in Uniprot KB, was neutralized by 4 μM varespladib, but not by marimastat, dimercaprol or DMPS. The SL1_ECHOC, SL124_ECHOC, VM3E2_ECHOC and VM3E6_ECHOC procoagulant candidates were not fully neutralized by varespladib and dimercaprol, but were effectively neutralized by 0.16 μM marimastat (i.e. a very low concentration) and by 20 μM DMPS. It could unfortunately not be determined which exact toxins were inhibited by varespladib and/or by dimercaprol, and to what proportion per toxin, as in these cases several venom toxins closely co-eluted. However, it is highly likely that the SVMPs are the dominant procoagulants given that it is known that SVMPs are procoagulant in this species and the SVMP inhibitors can block this activity. The PA2B8_DABRR, PA2B5_DABRR and PA2B3_DABRR were neutralized by 20 μM varespladib, but not influenced by marimastat and dimercaprol. Of these, only PA2B8_DABRR was reported to exhibit anticoagulant activity [2]. Only SLA_BITAR and SLB_BITAR were found back in the anticoagulation area (negative peak at retention time 16.7-17.1 min), which was fully inhibited by a low concentration of varespladib (0.8 μM). This anticoagulation area was not influenced by marimastat.

References

1.	Kemparaju, K.; Krishnakanth, T.; Gowda, T.V. Purification and characterization of a platelet aggregation inhibitor acidic phospholipase A_2_ from Indian saw-scaled viper (*Echis carinatus*) venom. *Toxicon* **1999**, *37*, 1659-1671.

2.	Faure, G.; Gowda, V.T.; Maroun, R.C. Characterization of a human coagulation factor Xa-binding site on *Viperidae* snake venom phospholipases A_2_ by affinity binding studies and molecular bioinformatics. *BMC structural biology* **2007**, *7*, 82.
